# A conserved NLR activation switch governs global transcriptional control of antibiotic biosynthesis in *Streptomyces*

**DOI:** 10.64898/2026.09.02.748808

**Authors:** Max L. Jordan, Hongrui Wang, Natalia M. Vior, Matthew J. Bush, Govind Chandra, Kathryn J. Stratton, Abbas Maqbool, Andrew W. Truman, Susan Schlimpert

## Abstract

Nucleotide-binding and oligomerization domain-like receptors (NLRs) are conserved molecular switches known for their role in immunity. A central feature in eukaryotic NLRs is a conserved sequence motif, acting as a regulatory latch whose destabilization triggers NLR activation.

Here, we show that this activation switch is conserved in the functionally distinct bacterial NLR-related transcription factor AfsR, which controls antibiotic production in *Streptomyces*. Disruption of this regulatory switch generates an autoactive AfsR variant that bypasses upstream signals and drives widespread transcriptional and metabolic reprogramming, increasing antibiotic production and activating otherwise silent biosynthetic pathways. Comparative genomics reveals a large family of AfsR-like regulators distributed across Actinomycetota that retain the conserved activation motif, suggesting a substantial reservoir of unexplored regulatory diversity that could be harnessed to expand antibiotic discovery. Together, these findings reveal a conserved mechanism linking NLR activation to transcriptional regulation in bacteria and suggest that this regulatory principle extends beyond AfsR.

## Introduction

Nucleotide-binding and oligomerization domain-like receptors (NLRs) are intracellular molecular switches that regulate diverse cellular processes across plants, animals, fungi and bacteria ^1–5^. Although best known for their roles in immunity, bacterial NLR-related proteins also function in non-immune processes, including transcriptional regulation of carbohydrate metabolism and secondary metabolite biosynthesis ^6–8^. However, how NLR-related proteins control these diverse activities outside canonical immune signaling remains poorly understood.

NLRs share a conserved tripartite architecture consisting of a variable N-terminal executioner domain, a central nucleotide-binding and oligomerization domain (NOD), and a C-terminal sensor domain composed of superstructure-forming repeats. The NOD contains a STAND-family P-loop NTPase, represented by NB-ARC domains in plants and NACHT domains in animals ^9,10^. Mechanistically, NLRs function as molecular switches in which nucleotide-dependent conformational changes control downstream activity via the N-terminal executioner domain ^11–16^. Central to this molecular switch is a conserved motif in the NB-ARC domain that locks the protein in an ADP-bound, autoinhibited state until effector-triggered nucleotide exchange induces the active conformation ^17–22^.

Beyond their well-established roles in innate immunity, STAND-family proteins also serve regulatory functions in bacteria. A notable example is found in the genus *Streptomyces*. These soil-dwelling bacteria are renowned for their complex multicellular life cycle and extraordinary capacity for specialized metabolism, which provides a rich source of bioactive molecules with applications in medicine and industry, including approximately two-thirds of clinically used antibiotics ^23–25^. Biosynthesis of these valuable molecules is governed by complex regulatory networks that include members of the *Streptomyces* antibiotic regulatory protein (SARP) family of transcriptional activators ^26^. A prominent example is the global regulator AfsR, which stimulates the production of diverse antimicrobial and antifungal metabolites ^27,28^.

Importantly, AfsR exhibits an NLR-type domain organization comprising an N-terminal SARP-family regulatory signaling module, a central NOD module containing an NB-ARC-type STAND ATPase, and a C-terminal tetratricopeptide repeat (TPR) domain (**Fig. 1A**). Thus, the SARP module provides the downstream regulatory output analogous to the N-terminal executioner domain of NLRs, while the C-terminal TPR domain is analogous to the repeat-containing sensor domain. AfsR binds a 22-bp repeat sequence in the promoter of *afsS*, a small regulatory gene whose expression promotes the biosynthesis of actinorhodin and undecylprodigiosin ^29^. Structural and biochemical studies have shown that *Streptomyces coelicolor* AfsR dimerizes via its SARP domain at the *afsS* promoter and recruits RNA polymerase to activate transcription ^29–31^. Additionally, AfsR activity depends on an intact NB-ARC domain and may be further modulated by phosphorylation and nutritional stress ^29,31^. However, whether AfsR employs the same conserved NLR activation switch as eukaryotic NLRs, and whether this switch could be directly engineered to control antibiotic biosynthesis independently of the natural upstream signal, remained unknown.

**Fig. 1.**
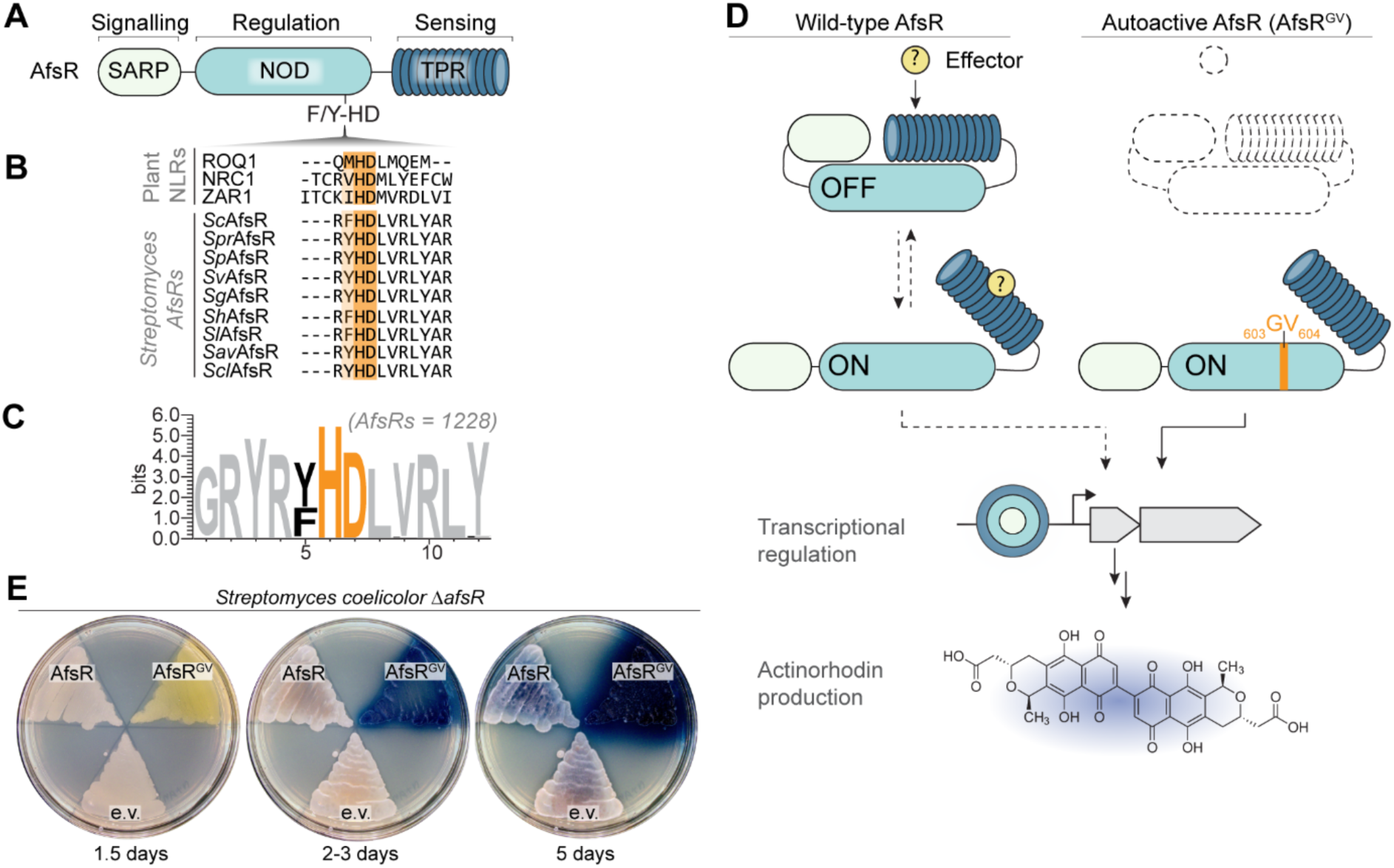
A conserved NLR activation switch controls AfsR-mediated antibiotic production in *Streptomyces coelicolor*. **(A)** Domain architecture of AfsR. SARP, *Streptomyces* Antibiotic Regulatory Protein; NOD, nucleotide-binding and oligomerization domain; TPR, tetratricopeptide repeats. **(B)** Multiple sequence alignment of the NOD module region encompassing the ‘MHD’ motif found in representative plant NLRs and published AfsR orthologs of *Streptomyces* species. Conserved residues are highlighted in orange. The UniProt identifiers of the NLR and AfsR orthologs shown in the alignment are: *Nicotiana benthamiana* ROQ1 (A0A290U7C4), *Solanum lycospersicum* NRC1 (A1X877), *Arabidopsis thaliana* ZAR1 (Q38834); *Streptomyces* AfsRs: *S. coelicolor* ScAfsR (P25941), S. *pristinaespirilis* SprAfsR (B5HE57), *S. peucetius* SpAfsR (Q4H0Z2), *S. venezuelae* SvAfsR (A5Z1X4), *S*. *griseus* SgAfsR (B1W547)*, S. hygroscopicus* ShAfsR (H2JUC6), *S. lividans* SlAfsR (A0A7U9DWY1)*, S. avermitilis* SavAfsR (Q82GU2), *S. clavuligerus* SclAfsR (E2Q034). **(C)** Sequence logo showing the conservation of the “MHD” motif in AfsR orthologs from *Streptomyces* (n=1228). **(D)** Cartoon detailing the effector-dependent switch from inactive (OFF) to active (ON) wild-type AfsR compared to the autoactive AfsR variant (AfsR^GV^). Activation of AfsR results in the production of the blue-pigmented antibiotic actinorhodin. **(E)** Actinorhodin production (blue) of *S. coelicolor* Δ*afsR* cells complemented with *afsR-3xFLAG*, *afsR^GV^-3xFLAG* expressed from its native promoter, or the empty vector (e.v.). Strains were grown on Difco nutrient agar and imaged at the indicated time points. Shown are representative images of duplicate experiments.

Here we show that a conserved sequence motif in the NOD module of the transcriptional regulator AfsR defines a universal activation switch that can be directly engineered to overcome the requirement for its native activating signal. By genetically locking AfsR into a constitutive ‘ON’ state, we bypass the need for complex upstream signaling cues, resulting in global metabolic reprogramming and enhanced antibiotic biosynthesis. Through integrated global genomics, *in vitro* DNA-binding assays, metabolomics, and comparative genomics, our work provides fundamental insights into the functional diversification of NLR-related proteins and demonstrates how AfsR autoactivation can be leveraged to increase specialized metabolite production and activate otherwise silent biosynthetic pathways under laboratory conditions in *Streptomyces*.

## Results

### A conserved activation switch in the bacterial NLR AfsR governs antibiotic biosynthesis in *Streptomyces*

In the absence of an effector, NLRs are maintained in an autoinhibited state by a network of intradomain interactions stabilized by bound ADP ^3^. A central governor of this inactive conformation is the highly conserved methionine-histidine-aspartate (MHD) motif within the nucleotide-binding and oligomerization domain. Several studies on plant NLRs have revealed that the histidine residue forms a critical hydrogen bond with the *β*-phosphate of ADP, effectively locking the NLR in an inactive conformation. Disrupting this intramolecular interaction, either through effector-triggered repositioning of the MHD motif or targeted mutagenesis, facilitates ADP-to-ATP exchange and triggers NLR activation ^17–22^.

Given the evolutionary relationship with plant NLRs ^9^, we hypothesized that AfsR may employ an analogous autoinhibitory mechanism. Sequence alignments of AfsR homologs and well-characterized representatives of plant NLRs revealed that the MHD motif is indeed conserved as an F/Y-HD (phenylalanine/tyrosine-histidine-aspartic acid) motif in *Streptomyces species* (n=1228) (**Fig. 1A-B, Supplementary Data 1**). AlphaFold3 modelling supports this homology, placing the histidine residue within the F/Y-HD motif of *S. coelicolor* AfsR (SCO4426, AfsR hereafter) in a structurally similar position to the regulatory histidine in the cryo-EM structure of the plant NLR ZAR1 ^21^ where it resides within the nucleotide-binding pocket, contacting the ADP moiety (**Fig. S1**). Consequently, we reasoned that disrupting the F/Y-HD motif in AfsR by site-directed mutagenesis would bypass the requirement for its unknown native effector, thereby converting AfsR into a constitutively active transcriptional regulator capable of driving antibiotic production regardless of the required stimulus (**Fig. 1D**).

To test this hypothesis, we engineered an AfsR variant carrying a histidine-to-glycine substitution at position 603 and an aspartic acid-to-valine substitution at position 604 (AfsR^GV^ hereafter). These specific changes have previously been shown to trigger autoactivation in the tomato NLR NRC1 ^19^. The *afsR* variant allele, expressed from its native promoter, was integrated *in trans* into the chromosome of an *S. coelicolor* Δ*afsR* variant. In parallel, we generated control strains carrying either wild-type *afsR* or the empty vector (e.v.) and confirmed the production of the AfsR variants by western blot analysis (**Fig. S2A**).

In *S. coelicolor*, AfsR positively regulates the production of actinorhodin, a characteristic, blue-pigmented antibiotic which is secreted into the surrounding medium. To assess the consequences of AfsR activation, *S. coelicolor* strains were grown on nutrient agar and imaged over five days. While cells expressing wild-type *afsR* and the empty vector control displayed low levels of actinorhodin production, cells producing the AfsR^GV^ variant showed visible production of, first, a yellow-pigmented molecule indicative of enhanced production of the otherwise silent polyketide coelimycin P1 ^33^, followed by a significant accumulation of a dark blue compound corresponding to actinorhodin (**Fig. 1E**). Spectrophotometric quantification of actinorhodin production in liquid-grown strains further supports these phenotypic observations (**Fig. S2B**), demonstrating that mutagenizing the conserved HD motif within the nucleotide-binding and oligomerization domain of AfsR results in a constitutively active AfsR^GV^ variant that can significantly induce actinorhodin production under both solid and liquid cultivation conditions.

### Autoactivation expands the regulatory activity of AfsR

To better understand the molecular basis for the activity of the AfsR^GV^ variant, we performed global transcriptomic analyses to compare gene expression profiles in *S. coelicolor cells* expressing either wild-type *afsR* or autoactive *afsR^GV^*. For this and all subsequent experiments, we used the standard growth medium YT, which supports dispersed growth of *Streptomyces* in liquid culture.

RNA-sequencing analysis revealed that cells producing autoactive AfsR^GV^ showed significant upregulation of antibiotic-associated genes, including those involved in the biosynthesis of actinorhodin, undecylprodigiosin, and coelimycin P1. These transcriptional changes were more pronounced in actively growing cells (2-day samples, **Fig. 2A-B**) compared to cells sampled at 5 days (**Fig. S3A-C** and **Supplementary Data 2**). Notably, we did not observe an effect on the transcript levels of *afsR* and *afsR^GV^*, indicating that the enhanced actinorhodin production associated with AfsR^GV^ is not attributable to differences in expression but likely reflects altered regulatory capacity. Among the most upregulated genes were two genes encoding transcriptional regulators: *afsS*, a known target of AfsR ^29^, and *SCO6715*, which encodes the largely uncharacterized WhiB-like regulator WblH ^34,35^ (**Fig. 2A** and **Fig. S3B**).

**Fig. 2.**
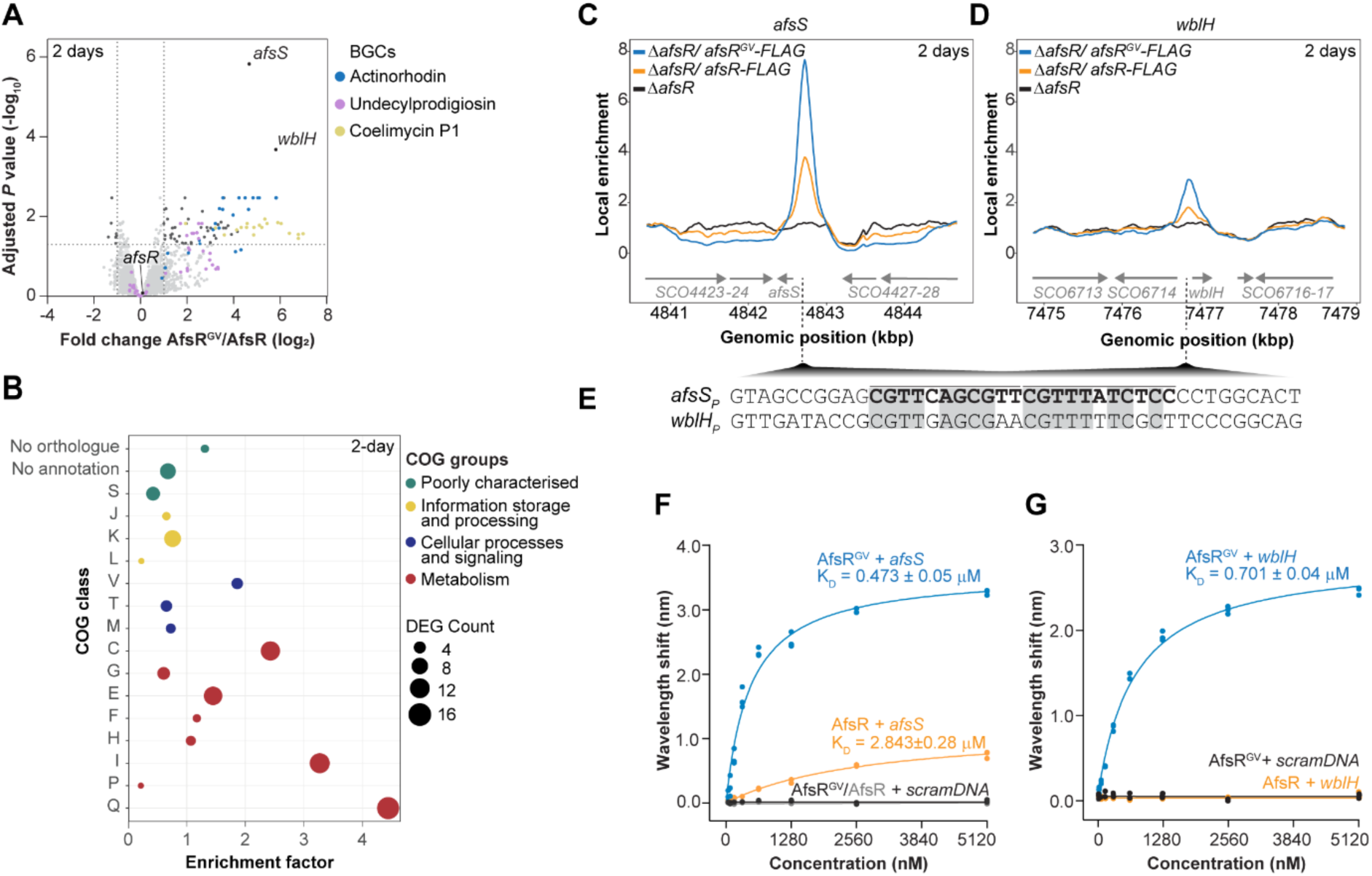
Autoactivation of AfsR drives global transcriptional reprogramming and alters DNA-binding specificity. **(A)** Volcano plot of differentially expressed genes (DEGs) in exponentially growing (2-day samples) *S. coelicolor* Δ*afsR* cells complemented with *afsR^GV^*compared to Δ*afsR* cells complemented with *afsR*. Data derived from 3 biological replicates. Dashed lines indicate significance thresholds. Selected genes and gene groups are identified by labels. **(B)** Analysis of Clusters of Orthologous Groups of proteins (COG) enriched in cells expressing *afsR^GV^* compared to *afsR* (2-day samples). Bubble size represents the number of DEGs assigned to each COG class. See Methods section for COG class definition. **(C)** and **(D)** ChIP-seq profiles showing AfsR^GV^-FLAG and AfsR-FLAG enrichment at the *S. coelicolor afsS* (C) and *wblH* (D) promoter region at 2 days. Analysis was performed on exponentially growing cells and 2 biological replicates. See also **Fig. S3A** and **S4C** (5 days). **(E)** Predicted 22nt DNA-binding sequence of AfsR^GV^ in the *wblH* promoter compared to the known AfsR binding motif in the *afsS* promoter (bold letters); conserved residues are shaded in grey. **(F)** and **(G)** Biolayer interferometry analysis of the interaction of increasing concentrations of purified AfsR^GV^ and AfsR with a double-stranded DNA fragment containing the putative binding sequence in the *afsS* (F) and *wblH* (G) promoter or scrambled DNA sequence (scramDNA). Experiments were performed in triplicate, and K_D_ values were determined by curve fitting. See also **Fig. S7**.

To define the genome-wide binding profiles of AfsR and its constitutively active variant, AfsR^GV^, we performed anti-FLAG chromatin immunoprecipitation followed by deep sequencing (ChIP-seq) using the same cultivation conditions and sampling time points used for the RNA-seq analysis (**Fig. S3A**). For this purpose, the *S. coelicolor* Δ*afsR* mutant was complemented *in trans* with a functional *afsR-FLAG* or *afsR^GV^-FLAG* allele (**Fig. S4A-B**). A non-FLAG-tagged Δ*afsR* strain was used as a negative control to eliminate background signals arising from non-specific antibody binding. Applying a twofold enrichment threshold, we identified five significantly enriched regions for AfsR-FLAG and three for AfsR^GV^-FLAG (**Supplementary Data 3**).

In the AfsR-FLAG dataset, one enriched region mapped to the *afsS* promoter at day 2 (**Fig. 2C** and **Fig. S4C)**. This region contains a conserved 22-bp AfsR-binding motif (**Fig. 2E**), corresponding to the previously reported AfsR-binding site ^29–31^, confirming the ability of the ChIP-seq approach to capture bona fide AfsR–DNA interactions. The remaining four AfsR-FLAG-enriched regions lacked the conserved AfsR-binding motif and were not associated with differential expression of associated genes (**Fig. 2A** and **Supplementary Data 2**), providing no additional evidence for direct transcription regulation under the conditions tested. One enriched region (*SCO4848*) was detected in both AfsR-FLAG and AfsR^GV^-FLAG datasets at both time points. Because this locus corresponds to the chromosomal integration site of the complementation constructs, we considered it an integration-associated artefact.

In contrast, the two remaining enrichment sites detected for AfsR^GV^-FLAG mapped to promoter regions upstream of *afsS* and *wblH* (*SCO6715*) (**Fig. 2C–D, Fig. S4C** and **Supplementary Data 3**). In both cases, RNA-seq analysis revealed significant differential expression of the corresponding genes (**Fig. 2A**), consistent with direct transcriptional regulation. While AfsR^GV^ occupancy at the *afsS* promoter was consistent with the previously characterized AfsR-dependent regulation of *afsS*, enrichment upstream of *wblH* represents a previously unidentified AfsR-binding site (**Fig.2D**). Motif analysis of the sequences underlying the ChIP-seq peak summits revealed a repeat element in the *wblH* promoter that closely resembles the characterized 22-bp AfsR-binding motif upstream of *afsS* ^29–31^ (**Fig. 2E**). This shared sequence feature supports a common mode of DNA recognition and is consistent with direct regulation of *wblH* by AfsR^GV^.

To corroborate the ChIP-seq results, we performed *in vitro* protein-DNA binding assays using Bio-Layer Interferometry. To this end, we used purified AfsR and AfsR^GV^ (**Fig. S5A-B**) and 42-bp double-stranded biotinylated DNA segments containing the predicted repeat sequences. In agreement with previous studies ^29–31^, we observed that wild-type AfsR bound the *afsS* promoter (KD = 2.843 ± 0.28 μM). The autoactive variant, AfsR^GV^, also recognized the *afsS* promoter sequence, but with a roughly five-fold higher affinity (KD = 0.473 ± 0.05 μM) (**Fig. 2F** and **Fig. S6**). Additionally, AfsR^GV^ bound the *wblH* promoter (KD = 0.701 ± 0.04 μM), whereas no binding was detected with the wild-type protein at this site (**Fig. 2G and Fig. S6**). Neither protein interacted with a scrambled control DNA sequence of the same length, confirming that the observed binding events are specific (**Fig. 2F-G** and **Fig. S6**).

Integration of *in vitro* DNA binding, RNA-seq, and ChIP-seq data indicates that autoactivating mutations expand the regulatory scope of AfsR. Therefore, AfsR autoactivation appears to bypass environmental or intracellular constraints that normally restrict promoter binding by the wild-type protein. As a result, in *S. coelicolor*, AfsR^GV^ drives increased expression of biosynthetic genes, likely mediated largely through enhanced transcription of its downstream targets *afsS* and *wblH*, which are widely co-conserved with *afsR* across *Streptomyces* species (**Fig. S7** and **Supplementary Data 1**).

### Autoactive AfsR can be used to induce the overproduction of specialized metabolites in *Streptomyces* species

To assess the global impact of autoactive AfsR^GV^ on specialized metabolite production in *S. coelicolor*, we performed untargeted liquid chromatography-mass spectrometry (LC-MS)-based metabolomic profiling. We analyzed whole culture extracts from Δ*afsR* strains expressing either wild-type *afsR* or autoactive *afsR*^GV^, obtained after two and five days of fermentation (**Fig. S3A** and **Supplementary Data 7**). Principal component analysis revealed a profound and distinct separation between the metabolic profiles of cells expressing autoactive *afsR^GV^* and wild-type *afsR* in five-day samples (**Fig. S8A**). This point marked the transition to the stationary growth phase in the wild type, a phase generally associated with enhanced metabolite accumulation in streptomycetes. We therefore focused subsequent analyses on the five-day samples. Comparative metabolomic analysis corroborated the global divergence of metabolite profiles in cells expressing *afsR^GV^* (**Fig. 3A**), showing significant differences in the abundance of hundreds of metabolites, consistent with the effects of a global regulator such as AfsR. A targeted search for well-characterized metabolites from *S. coelicolor* confirmed the overproduction of actinorhodin, undecylprodigiosin and coelimycin P1-related molecules in response to autoactive AfsR^GV^ (**Fig. 3B**, **Fig. S8B** and **Supplementary Data 7-8**). These results are further supported by our RNA-seq data, showing transcriptional upregulation of the corresponding BGCs (**Fig. 2A**). Notably, the highlighted metabolites account for only a fraction of the metabolic changes observed in *afsR^GV^*-expressing cells. Using mass-spectral molecular networking, we found that autoactive AfsR^GV^ induces the production of diverse and unique metabolites of largely unknown identity (**Fig. 3C**).

**Fig. 3.**
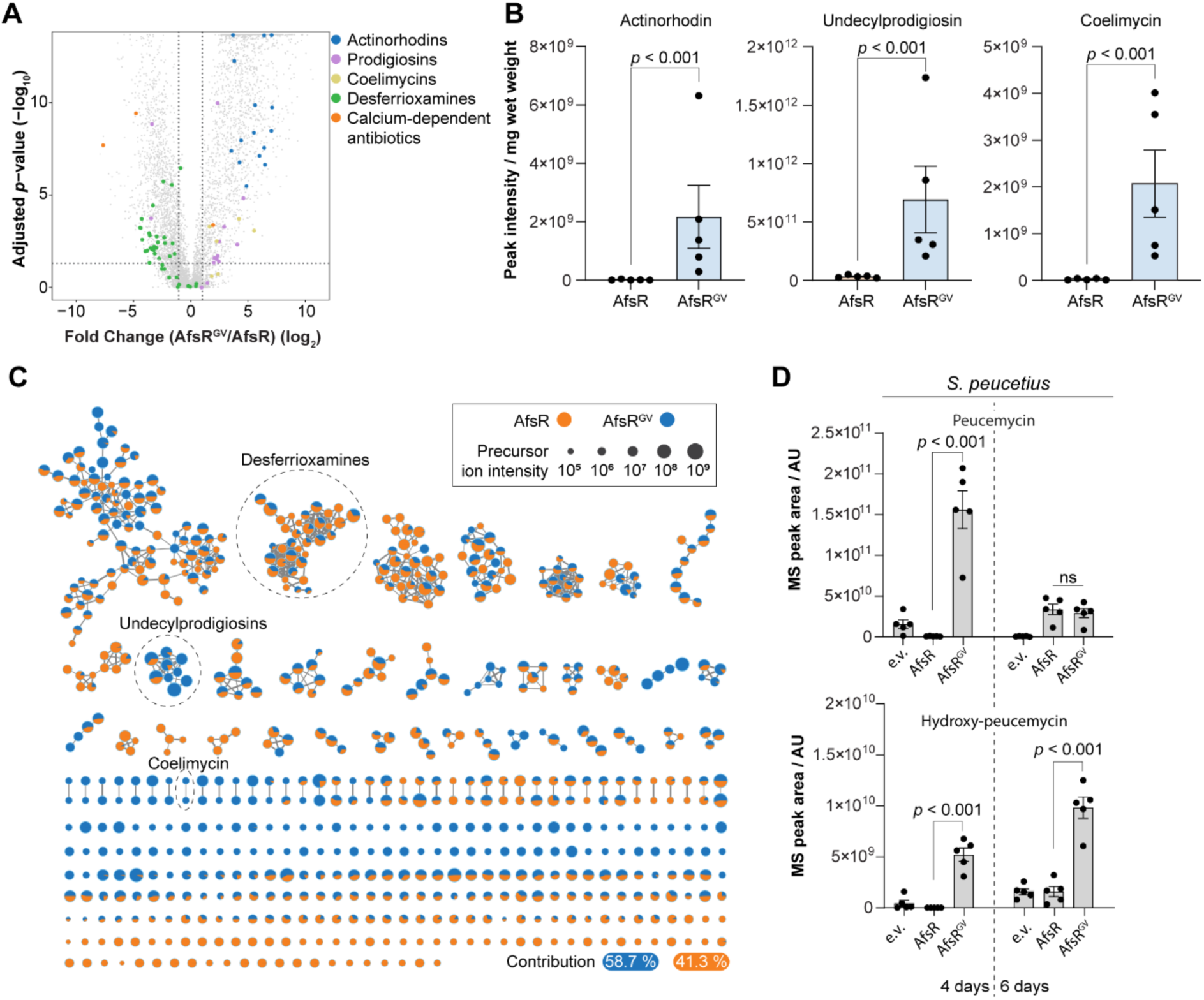
Autoactivation of AfsR induces a global metabolomic response in *Streptomyces*. **(A)** Volcano plot of differentially produced metabolite features detected by LC-MS in crude extracts of *S. coelicolor* Δ*afsR* expressing *afsR^G^* compared to cells expressing *afsR*. Dashed lines indicate significance thresholds. Known metabolite features are identified by labels. **(B)** Bar charts comparing the production of selected metabolites in *S. coelicolor* Δ*afsR* expressing *afsR* or *afsR^GV^* at 5 days of fermentation. The MS peak area was normalized by the cell pellet weight for each sample. Error bars represent the standard error (n=5). Statistical significance (*P* value) was determined using the Wilcoxon rank sum test. **(C)** Molecular networks analysis of culture extracts from *S. coelicolor*. Nodes represent metabolites, with sizes proportional to precursor ion intensity, and pie charts indicate relative abundance in cells expressing *afsR* (orange) versus *afsR^GV^* (blue). Edges are weighted by MS/MS cosine similarity scores. Dashed circles indicate groups of known molecules. **(A-C)** Data derived from 5 days of fermentation in YT broth, using 5 biological replicates. **(D)** Bar charts comparing the production of peucemycin and hydroxy-peucemycin detected in crude extracts of *S. peucetius* expressing *afsR, afsR^GV^* from the native promoter. e.v. is an empty vector control. MS Peak areas are normalized by culture growth (colorimetric DNA measurement). Data are derived from 4 and 6 days of fermentation in SFM broth, with 5 biological replicates per strain. Error bars represent the standard error. Statistical significance (adjusted *P* value) was determined using Kruskal-Wallis tests followed by a Dunn test with BH correction for peucemycin and one-way ANOVA followed by Turkey’s HSD post-hoc test for hydroxy-peucemycin; ns, not significant. See also Fig. **S10**.

To evaluate whether AfsR autoactivation represents a transferable strategy for unlocking biosynthetic potential across the genus, we applied this approach to *Streptomyces peucetius* var. *caesius* (hereafter *S. peucetius*), the native producer of the anthracycline antitumor agents daunorubicin and doxorubicin ^36^. In addition to the well-understood anthracycline pathway, the *S. peucetius* genome is predicted to encode 24 BGCs, many of which remain uncharacterized ^37,38^. *S. peucetius* AfsR (Sp-AfsR) shares 70.8% sequence identity with *S. coelicolor* AfsR and retains the conserved histidine-aspartic acid residues within the MHD motif (**Fig. 1B**), suggesting that the autoactivation mechanism may be conserved. Using a strategy analogous to that established in *S. coelicolor*, wild-type *Sp-afsR* and the autoactive variant allele *Sp-afsR^GV^*, encoding histidine-to-glycine and aspartic acid-to-valine substitutions at positions 594 and 595, respectively, were integrated *in trans* into the chromosome of wild-type *S. peucetius* and expressed from the native promoter.

Using LC-MS, we analyzed extracts of culture supernatants collected after four and six days of fermentation. Under these conditions, we did not detect doxorubicin production and did not observe significant daunorubicin production in cells expressing *afsR^GV^* compared with the control strains (**Fig. S8C**). However, as observed for *S. coelicolor*, expression of autoactive Sp-*afsR^GV^* also affected the global metabolome of *S. peucetius*, resulting in the production of multiple distinct families of molecules compared to cells expressing wild-type Sp-*afsR* or the empty vector control (**Fig. S9** and **Supplementary Data 9**). While the vast majority of the detected metabolites remain unknown, one of the most enriched networks in cells expressing Sp-*afsR^GV^*, corresponds to peucemycin-related molecules (**Fig. 3D** and **Fig. S10**). Peucemycins are a group of recently discovered specialized metabolites that display antibacterial and anticancer activity ^39^. Importantly, the corresponding biosynthetic gene cluster is silent under standard laboratory conditions ^39,40^. Collectively, our results show that utilizing a constitutively active AfsR^GV^ variant can trigger the expression of silent BGCs and significantly enhance the production of known metabolites with clinically relevant bioactivities.

### Actinomycetes encode multiple AfsR-like regulators with unknown regulatory targets

SARP-type regulators, such as AfsR, are widespread in *Streptomyces* and play a central role in regulating the expression of BGCs, making this group of regulators attractive targets for genetic engineering approaches to facilitate the discovery of novel bioactive compounds ^26^. SARPs are classified primarily by their size and domain complexity into small, medium, and large classes ^26,28^. AfsR is a typical member of the “large SARPs”, which can be further split into two structural subgroups. The first subgroup contains regulators that share the same characteristic tripartite domain architecture as AfsR: an N-terminal SARP domain, a central nucleotide-binding and oligomerization domain of the NB-ARC clade, and a series of C-terminal tetratricopeptide repeats. The second subgroup includes SARP-LALs (Large ATP-binding regulators of the LuxR family) proteins, which lack tetratricopeptide repeats and instead carry a C-terminal LuxR domain. While previous analyses have highlighted the distribution and regulatory potential of LuxR-type, small and medium-sized SARPs in *Streptomyces*, much less is known about members of the large SARP-type regulator family that are structurally similar to AfsR.

To determine the abundance of AfsR-like regulators in *Streptomyces* species, we first performed a relaxed reciprocal BLAST search using the AfsR sequence against the 1,228 *Streptomyces* genomes previously identified as encoding a reciprocal best BLAST hit for AfsR (**Supplementary Data 4**). We then applied a series of filtering steps to remove small and medium-sized SARPs, and used InterProScan ^41^ to exclude proteins lacking any of the characteristic AfsR functional domains. This pipeline identified a total of 2670 AfsR-like regulators (**Fig. 4A**). While some *Streptomyces* genomes (n=197) only encode the primary AfsR ortholog, the majority of the genus is predicted to harbor between 1 and 4 additional AfsR-like regulators (n=924). A small subset of genomes (n=107) encodes even higher numbers, with some carrying up to 11 additional AfsR-like regulators (**Supplementary Data 4**), further pointing to the underlying complexity in the regulation of specialized metabolite production across the genus.

**Fig. 4.**
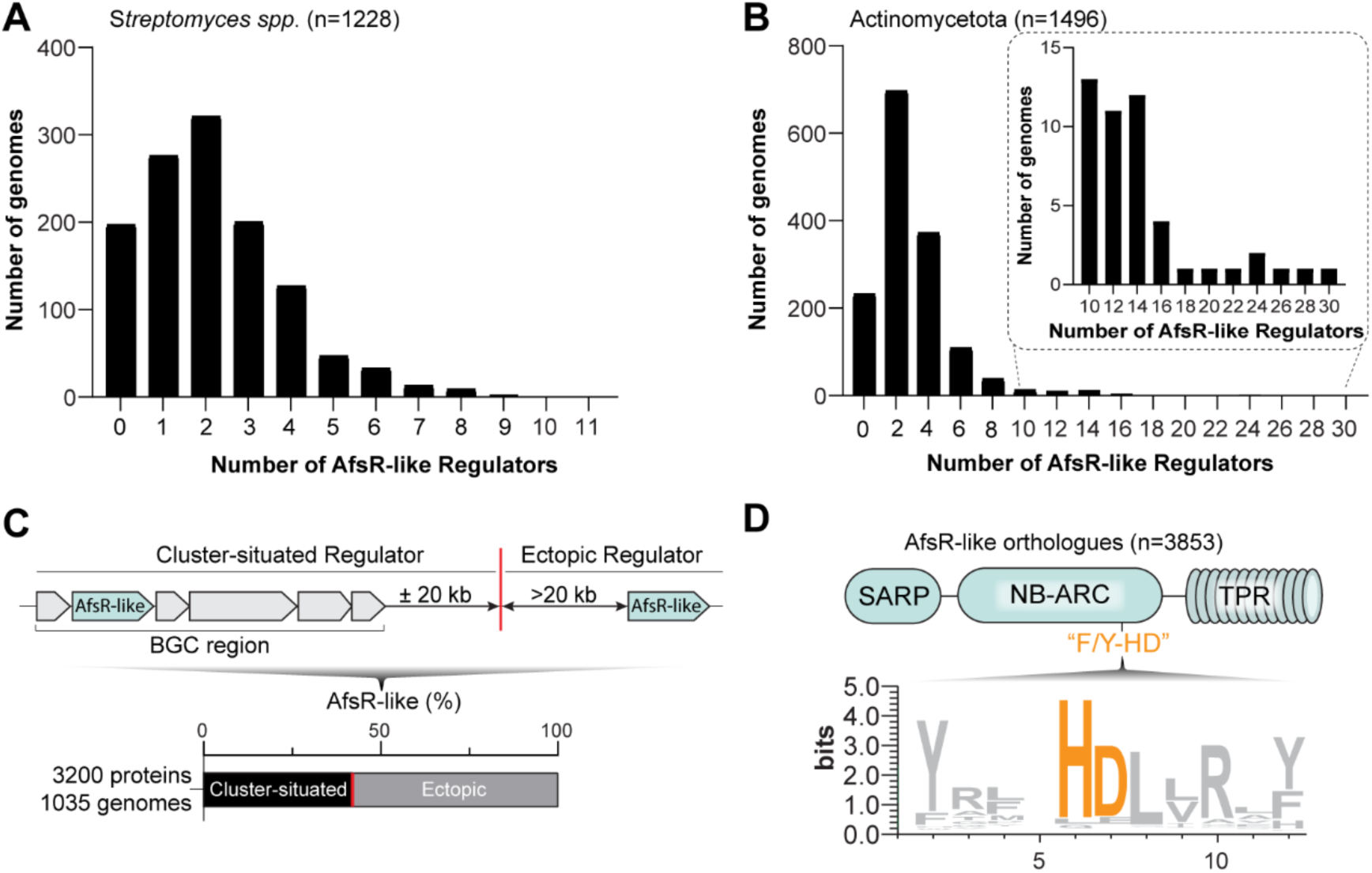
Actinomycetota encode additional, largely uncharacterized AfsR-like regulators. **(A)** Distribution of additional AfsR-like proteins (n=2670) across *Streptomyces* genomes (n=1228). **(B)** Total number of AfsR-like regulators (n=3853) detected in Actinomycetota genomes (n=1496, including *Streptomyces*) that encode AfsR. **(C)** Ratio of AfsR-like proteins (n=3200) across Actinomycetota that can be classified as “Cluster-situated” or “Ectopic” regulators based on their genomic association with biosynthetic regions predicted by antiSMASH. **(D)** Sequence logo showing the conservation of the F/Y-HD motif in identified Actinomycete AfsR-like regulators (n=3853).

To assess whether this regulatory expansion is a unique feature of *Streptomyces* or a broader characteristic of specialized metabolite producers, we expanded our phylogenomic search across the wider Actinomycetota phylum. Screening 4,500 Actinomycetota genomes, we identified a primary AfsR ortholog in 1,496 genomes (**Supplementary Data 4**). Consistent with previous reports, the vast majority of these (n=1,228, 82%) were restricted to the genus *Streptomyces* ^26,28,42^. Across the entire phylum dataset, we identified 3,853 AfsR-like regulators (**Fig. 4B**), including a particular enrichment in several rare actinomycetes such as *Kutzneria*, *Lentzea*, *Saccharothrix* and *Amycolatoposis* ^43^ (**Supplementary Data 4**).

To better understand the functional context of these regulators, we explored their genomic association with biosynthetic regions predicted in the antiSMASH database ^37^. This analysis was therefore restricted to the 3,200 AfsR-like proteins identified in genomes with available antiSMASH database annotations, representing 69% of the total genomic dataset and 83% of the identified AfsR-like proteins. Regulators were classified as "cluster-situated" if they were encoded within a predicted BGC boundary or within a 20 kb flanking region (upstream or downstream). Regulators located outside of these boundaries were classified as "ectopic". Our analysis revealed that 41.66% (n = 1,333) of the AfsR-like proteins are encoded within or adjacent to BGCs, while the remaining 58.34% (n = 1,867) reside in ectopic locations and appear not to be associated with specific BGCs, similar to AfsR (**Fig. 4C** and **Supplementary Data 5** and **6**), suggesting a substantial role for these proteins beyond local pathway-specific regulation.

Finally, we investigated whether the established synthetic activation approach for classic AfsR regulators could be universally applied to the newly identified AfsR-like proteins. To evaluate this, we aligned the nucleotide-binding and oligomerization domains of the 3,853 identified AfsR-like sequences to generate a conserved sequence motif. This analysis revealed that the critical "HD" motif is highly conserved across the entire expanded family of regulators (**Fig. 4D**). Since our data demonstrate that mutating the "HD" motif in AfsR results in its autoactivation (**Fig. 1**), the high level of conservation of these residues indicates that a similar genetic engineering strategy could be used to generate active variants of AfsR-like regulators. Extending this approach across the phylum could therefore help overcome native regulatory repression and elicit expression of otherwise silent BGCs.

## Discussion

NLRs function as conserved molecular switches, held inactive by a regulatory latch whose disruption triggers the transition to the active state. Our results show that this activation principle is not unique to eukaryotic immune receptors but is preserved in AfsR, a bacterial NLR transcription factor that governs specialized metabolite production in *Streptomyces*. Crucially, this activation switch can be directly harnessed to control antibiotic gene expression independently of native upstream signals, providing an experimental strategy to access the biosynthetic potential encoded within *Streptomyces* genomes and expand our understanding of the functional diversity of bacterial NLR proteins.

Like eukaryotic NLRs, AfsR activity appears to be controlled by a transition between inactive and active states. In NLRs that function as immune receptors, dysregulation of this activation process can lead to inappropriate immune signaling, fitness penalties and cell death ^44,45^. In contrast, our *in vivo* and *in vitro* analyses show that autoactivation of AfsR drives global transcriptional reprogramming through enhanced promoter recognition, resulting in a significant metabolic shift relative to the wild type. We propose that the autoactive AfsR^GV^ variant presents a constitutive mimic of the active, ‘ON’ state of AfsR. In canonical NLRs, transition to this state is controlled by the C-terminal sensor domain, which maintains the protein in an auto-inhibited ‘OFF’ state until a specific effector is detected ^15,46–48^. Conservation of the HD motif in AfsR enables direct engineering of this universal activation switch to bypass upstream regulatory control, without requiring prior identification of the natural activating signal. The HD-to-GV substitution within this motif (referred to as MHD in plant NLRs) likely disrupts the autoinhibited state, which may facilitate ADP-to-ATP exchange and promote conformational rearrangements that enhance the interaction of AfsR with its target promoters. This model is consistent with previous studies demonstrating that the NOD module and TPR sensor domain negatively regulate AfsR transcriptional activation, whereas ATP binding promotes AfsR activity and conformational changes ^29,31^. Furthermore, mutation of a conserved histidine residue involved in ADP binding, equivalent to AfsR^H603^, within the NB-ARC and NACHT domain of several plant and human NLRs, as well as in bacterial bNACHT antiphage systems, has been shown to result in constitutive activation, suggesting a conserved role for this residue in NLR autoinhibition across the tree of life ^4,19,49,50^.

Our *in vivo* and *in vitro* DNA-binding data support a model in which autoactivation of AfsR alters its DNA-binding activity. Specifically, significant binding to the *wblH* promoter was observed only with autoactive AfsR, whereas binding to the *afsS* promoter was enhanced approximately fivefold relative to wild-type AfsR. These results suggest that the *afsS* promoter is a high-affinity target that can be recognized through the intrinsic DNA-binding activity of AfsR, whereas interaction with less conserved repeat sequences, such as those present in the *wblH* promoter, requires the activated state. Thus, activation of AfsR may not simply increase DNA-binding affinity but also broaden its promoter recognition repertoire, enabling recruitment to lower-affinity regulatory sites. A possible explanation for the distinct DNA-binding behavior could be that activation induces a conformational rearrangement of AfsR that alters its interactions with DNA. This hypothesis would be compatible with two recent cryo-EM structures showing that wild-type AfsR activates transcription as a dimer in complex with the *afsS* promoter and RNA polymerase ^30,31^. However, our findings suggest that the current AfsR cryo-EM structures may represent only one functionally relevant state of active AfsR, while additional conformations that enable recognition of a broader range of promoters remain to be characterized.

The observed expansion of regulatory output by activated AfsR has direct metabolic consequences, as reflected in our metabolomic analyses of two *Streptomyces* species. Importantly, while *Streptomyces* species harbor a large repertoire of natural product BGCs, the vast majority of these gene clusters are poorly expressed under standard laboratory conditions ^51^. Here, we show that expression of autoactive *afsR* variants elicits overproduction of known molecules, such as actinorhodin and undecylprodigiosin, and triggers the expression of the silent peucemycin and coelimycin BGCs ^33,40^ as well as structurally diverse molecules of unknown identity. We note that expression of the autoactive *afsR* variant reduces growth rate, indicating a classic trade-off observed when metabolic flux is redirected to specialized metabolism ^52^. While developing a universal engineering strategy to awaken the biosynthesis of valuable specialized metabolites remains challenging, combining the expression of an autoactive *afsR* variant with targeted host metabolic engineering ^53^ provides a potential framework to simultaneously bypass upstream signal requirements and overcome primary metabolic bottlenecks.

Finally, our comparative genomic analyses reveal an expanded family of AfsR-like regulators that extend beyond the genus *Streptomyces* and that reside both within and outside predicted BGCs. These results underscore the multi-level transcriptional regulation of specialized metabolism in Actinomycetota and support the notion that an interplay between global and BGC-associated regulators enables simultaneous or sequential cross-regulation of multiple biosynthetic pathways ^54^. Crucially, the conservation of the HD motif within the NB-ARC domains of various AfsR-like proteins suggests that this locus serves as a universal regulatory checkpoint. Leveraging the principles validated in this study, targeted engineering of this specific pocket provides a transferable strategy to bypass upstream signaling and induce autoactivation in homologous regulators throughout the phylum Actinomycetota. Importantly, this approach does not require prior knowledge of the natural effector for each individual regulator because the universality of the activation switch, rather than the identity of the upstream signal, enables the strategy to scale across the phylum.

In conclusion, by demonstrating that the regulatory principles of eukaryotic immune receptors can be rationally engineered to control bacterial transcription, this work broadens our understanding of the functional diversity and evolutionary versatility of the NLR protein family. Here, we have shown that the NLR-related transcription factor AfsR operates as a tightly regulated molecular switch whose autoinhibitory constraints can be bypassed to access specialized metabolism in *Streptomyces* and related genera. Thus, the described AfsR autoactivation strategy provides an alternative framework for awakening silent BGCs, offering a powerful addition to the synthetic biology toolkit for natural product discovery and metabolic engineering in other biological systems that employ NLR-type regulators.

## Materials and Methods

### Strain and Plasmid Construction

All strains used in this work are listed in **Supplementary Table S1**. Details of the plasmids and a list of oligonucleotides used in this study are provided in **Supplementary Tables S2** and **S3**, respectively.

Plasmids were introduced into *S. coelicolor* via intergenic conjugation as previously described ^55^. For *S. peucetius*, intergeneric conjugation was performed using fragmented liquid-grown mycelia instead of spores. Recipient cultures were grown in 30 mL tryptic soy broth (TSB) until confluent. Mycelia were harvested by centrifugation at 1,127 × *g*, resuspended in 2 mL TSB, and mechanically disrupted by vortexing at maximum speed 10 times. The fragmented mycelia were then washed once with 30 mL of TSB and concentrated by resuspension in a final volume of 800 µL TSB. For each conjugation reaction, a 100 µL aliquot of this mycelial suspension was mixed with the donor strain. Conjugation mixtures were plated on solid Soy Flour Mannitol (SFM) medium; for *S. peucetius*, the medium was supplemented with 10 mM MgCl2.

### Media and Culture Conditions

*E. coli* strains were routinely grown in lysogeny broth (LB) medium and *S. coelicolor* and *S. peucetius* were maintained on solid Soy Flour Mannitol (SFM) media ^55^. For liquid cultures, *S. coelicolor* was routinely grown in a 50:50 mix of Yeast extract–Malt extract and tryptic soy broth (YT), while *S. peucetius* was grown in tryptic soy broth (TSB) or SFM. When appropriate, media were supplemented with the following antibiotics: hygromycin (25 µg/mL), apramycin (50 µg/mL), nalidixic acid (25 µg/mL), kanamycin (50 µg/mL), carbenicillin (100 µg/mL) and chloramphenicol (25 µg/mL).

### Identification of AfsR homologs in Actinomycetota

To identify AfsR homologs, a forward BLAST search was performed with the protein sequence of the *S. lividans* AfsR orthologue (SlAfsR, SLI_4664) using default parameters against a custom database of Actinomycete proteomes ^56,57^. This database was constructed using the bacterial assembly summary dataset downloaded from the NCBI GenBank FTP server on April 2, 2025. The dataset was filtered to include only Actinomycete ‘complete’ genomes assembled into fewer than 9 contigs with the corresponding protein FASTA files (_protein.faa.gz) available on the NCBI FTP server. These proteomes were downloaded, with the proteomes of *Streptomyces lividans* 1326 (GCA_000403665.1) and *Streptomyces coelicolor* A3(2) (GCA_000203835.1) manually integrated into the database to serve as internal references. The list of genomes encoding an AfsR homolog is provided in **Supplementary Data 1.**

Hits obtained from the initial forward BLAST search were used as queries for a reverse BLAST search against the *S. lividans* 1326 proteome database. Reciprocal best hits (RBHs) were identified by filtering for sequences that, based on bitscore, were the top forward BLAST hit for SlAfsR in that proteome during the forward search and had SlAfsR as the top reverse hit. To exclude smaller or medium-sized *Streptomyces* Antibiotic Regulatory Proteins (SARPs) (Augustijn et al., 2025; Yan and Xia, 2024), candidate sequences were filtered to retain only those with an aligned length greater than 800 amino acids. Conversely, potential multi-domain fusion artefacts or expanded proteins, which were defined as orthologs exceeding 1.25 times the mean sequence length, were manually inspected using the InterProScan web server. This manual curation identified two sequences containing extraneous domains (QPP05265.1 from GCA_015910445.1 and WXK67064.1 from GCA_045291505.1), which were excluded from subsequent analyses.

To definitively confirm the identity of the remaining candidate orthologs, local domain annotation was performed using InterProScan v5.52-86.0 ^41,58^ on the NBI High-Performance Computing (HPC) cluster. Sequences were validated based on the presence of the characteristic domains defined for AfsR and large SARPs ^26^: a C-terminal transcriptional regulatory protein domain (PF00486), a bacterial transcriptional activator domain (BTAD; PF03704), a P-loop NTPase (SSF52540), an NB-ARC domain (PF00931), and tetratricopeptide repeats (TPRs; G3DSA:1.25.40.10). To account for overlapping BTAD and TPR annotations, protein domain annotations were verified to ensure that a valid TPR annotation resided downstream of the P-loop NTPase domain.

### Identification of Actinomycete AfsR-like proteins

To identify broader distribution of AfsR-like regulators across the filtered Actinomycetota dataset, the reverse BLAST data was re-examined. True AfsR orthologs identified in the previous reciprocal best hit (RBH) pipeline were excluded from this analysis. Remaining sequences were filtered to retain only those with an aligned length greater than 500 amino acids, effectively removing small SARPs ^26,28^. Functional domain annotation was performed locally using InterProScan v5.52-86.0 on the NBI HPC cluster ^41,58^. Candidate sequences were required contain the characteristic domains previously defined for AfsR ^26^, and the total number of verified AfsR-like sequences was quantified per genome assembly.

To determine the genomic distribution of these AfsR-like regulators relative to biosynthetic gene clusters (BGCs), predicted BGC boundaries were retrieved from the antiSMASH database (version 5; accessed April 27, 2026) ^37^. Predicted BGC regions were filtered for the phylum Actinomycetota, and the ‘NCBI accession’ field cross-referenced to the encompassing RefSeq (GCF) assemblies. AfsR-like protein entries were mapped to their corresponding genomic positions by cross-referencing GenBank locus_tag identifiers with their respective RefSeq coordinates, using the ‘gbrs_paired_asm’ field of the assembly summary to facilitate matching.

Out of 1,264 genomes encoding a total of 3,853 identified AfsR-like proteins, 1,035 genomes (representing 3,200 AfsR-like proteins) were successfully cross-referenced with the antiSMASH database ^37^. To account for instances where BGC boundaries might extend beyond default algorithmic predictions, a flanking window of 20 kb was applied upstream and downstream of each predicted antiSMASH BGC boundary. AfsR-like coding genes were then classified and quantified based on whether they localized strictly within the predicted BGC borders, within the 20 kb flanking windows, or entirely outside of these biosynthetic boundaries.

### Identification of WblH and AfsS Orthologs

To identify WblH orthologs, a reciprocal BLAST analysis was performed as described above, utilizing the *S. lividans* 1326 WblH sequence (SLI_7059; EOY51763.1) as the initial query.

Due to the high sequence variability of AfsS, BLAST homology searches were insufficient. Instead, putative AfsS orthologs were identified based on genomic synteny. Using the corresponding GFF3 annotation files for each genome, the genomic regions immediately downstream of the previously identified *afsR* orthologs were screened for short open reading frames encoding proteins of fewer than 200 amino acids. To confirm their identity despite low primary sequence conservation, the resulting candidate sequences were analyzed using the RADAR (Rapid Automatic Detection and Alignment of Repeats) software ^59,60^ to verify the presence of the characteristic repeats intrinsic to AfsS orthologs ^29,61^. The list of identified WblH and AfsS orthologs is provided in **Supplementary Data 1.**

### Structural alignment of AfsR and ZAR1

A structural model of AfsR (SCO4426) with ADP was generated using the AlphaFold3 server ^62^. The auto seed option was used, resulting in seed 576924199. The resulting structure was aligned to the NOD domain (aa 145-514) of the experimentally determined ZAR1-ADP structure (PDB: 6J5W) using ChimeraX ^15,63^.

### Identification of Autoactivity-Associated Residues

The nucleotide-binding and oligomerization domain (NOD) was defined as the region spanning from the N-terminus of the P-loop NTPase domain (SSF52540) to the C-terminus preceding the superstructure-forming repeats consisting of Tetratricopeptide Repeats (TPRs; G3DSA:1.25.40.10) or Ribonuclease Inhibitor (G3DSA:3.80.10.10). For comparative analysis, NOD sequences were extracted from a reference set of verified AfsR orthologs from *S. coelicolor* (P25941), S. *pristinaespirilis* (B5HE57), *S. peucetius (*Q4H0Z2), *S. venezuelae* (A5Z1X4), *S*. *griseus* (B1W547)*, S. hygroscopicus* (H2JUC6), *S. lividans* (A0A7U9DWY1)*, S. avermitilis* (Q82GU2), *S. clavuligerus* (E2Q034) and the plant NLR immune receptors *Solanum lycopersicum* NRC1 (A1X877), *Nicotiana benthamiana* ROQ1 (A0A290U7C4) and *Arabidopsis thaliana* ZAR1 (Q38834). These sequences were aligned using MAFFT v7.520 ^64^ on the NBI HPC cluster, with iterative refinement (--maxiterate 1000) and local pairwise alignment (--localpair) constraints.

To investigate the conservation of the MHD motif within the actinomycete AfsR orthologs and AfsR-like proteins, the corresponding NOD sequences were extracted and aligned using MAFFT v7.520 under the automatic choice of strategy mode. The resulting multiple sequence alignments were trimmed using a custom Python script to isolate the region immediately flanking the predicted F/Y-HD motif of the *S. coelicolor* AfsR reference sequence. To visualize amino acid conservation and frequency across this motif, a sequence logo was generated from the trimmed alignment using WebLogo3 ^65,66^.

### Protein Extraction and Western Blotting

To prepare protein samples, 2 mL aliquots of liquid *S. coelicolor* YT cultures were sampled at the desired time points. Mycelium was pelleted by centrifugation at 4 °C, washed once with wash buffer (20 mM Tris-HCl pH 8.0, 5 mM EDTA), and snap-frozen in liquid nitrogen. Frozen cell pellets were thawed on ice, resuspended in ice-cold lysis buffer (20 mM Tris-HCl pH 8.0, 5 mM EDTA, 1x EDTA-free protease inhibitor cocktail; Sigma-Aldrich), and sonicated on ice at an amplitude of 4.5 µm for 9 cycles (15 s on/15 s off). The resulting cell lysates were cleared by centrifugation at maximum speed for 15 min at 4 °C. Total protein concentration in the cleared lysates was quantified using the Bradford reagent (Bio-Rad). Equivalent total protein concentrations (0.4 mg/mL) were resolved using 4-20% Tris-Glycine SDS-PAGE precast gels (Invitrogen) and analyzed via Western blotting using the following antibodies: rabbit anti-FLAG (1:5,000; Sigma, cat. #F7425), rabbit anti-AfsR (1:2,500; generated in this study), and goat anti-rabbit IgG HRP-conjugate (1:10,000; Abcam, cat. #ab6721).

### Quantification of Actinorhodin Production

To quantify total actinorhodin production in liquid-grown cultures, a spectrophotometric assay was performed as previously described ^55^. A 400 µL aliquot of each sample was mixed with 100 µL of 5 M KOH, thoroughly vortexed, and centrifuged at 3,000 x *g* for 5 min to pellet cellular debris. A 100 µL aliquot of the resulting supernatant was transferred to a clear 96-well microplate, and the absorbance was measured at 640 nm using a SPECTROstar Nano microplate reader (BMG LABTECH). To normalize the absorbance readings against biomass, the remaining sample mixture was centrifuged at 17.000 x *g* for 30 min, the supernatant was completely aspirated, and the wet cell pellet mass was recorded. Final actinorhodin values were expressed relative to the wet biomass weight.

### *S. coelicolor* LC-MS sample preparation

Metabolomics experiments were performed in biological quintuplicate. Cultivations were conducted in 250 mL Erlenmeyer flasks equipped with stainless steel spring coils to facilitate mycelial dispersion, containing 30 mL of YT medium, inoculated with approximately 10^6^ spores, and incubated at 28 °C with shaking at 250 rpm. After 2 and 5 days of incubation, 500 µL samples of each culture were withdrawn and stored at -20 °C until processing. For metabolite extraction, the samples were thawed on ice, and a 400 µL aliquot of each culture was mixed with an equal volume (400 µL) of 100% methanol. The mixtures were incubated at room temperature for 30 min with vigorous shaking at 1,200 rpm. Cellular debris and precipitated proteins were pelleted by centrifugation at 17,000 x *g* for 15 min. Finally, 600 µL of the clear supernatant was transferred to high-performance liquid chromatography (HPLC) vials for downstream analysis.

### *S. peucetius* LC-MS sample preparation

Seed cultures were grown in TSB media for 72 hours before being normalized to an OD_600_ of 1 and 1 mL of this culture was added to 30 mL SFM media in a sprung flask. At sampling points, 1 ml samples were taken and stored at -20 °C. Upon use, samples were thawed and centrifuged at 17,000 x *g*. The resulting supernatant was placed in HPLC vials for analysis. DNA was quantified in the sample using the Burton Assay to normalize metabolomics data ^67^.

### LC-MS and MS/MS data acquisition

High-resolution LC-MS and MS/MS data for both *S. coelicolor* and *S. peucetius* samples were obtained using a Hybrid Quadrupole-Orbitrap Q-Exactive mass spectrometer coupled to a Vanquish UHPLC system (Thermo Fisher Scientific). 5 μL of the *S. coelicolor* samples and 10 μL of *S. peucetius* samples were injected onto a Phenomenex Kinetex 2.6-µm C18 column (50 mm × 2.1 mm, 100 Å) set at a temperature of 40 °C. Mobile phases A (water + 0.1% (v/v) formic acid) and B (100% acetonitrile) were used in a 9.2-minute method consisting of a gradient from 5% to 95% B over 6 minutes, followed by a wash at 95% B and re-equilibration to 5% B, with a flow rate of 0.6 mL/min. Mass spectra were obtained in positive and negative mode for *S. coelicolor* and positive mode only for *S. peucetius* using full MS and data dependent MS2 (full MS/dd-MS2) with the following acquisition settings: chromatography peak width = 7s; Full MS settings: resolution = 70,000, AGC target = 3x10^6^, maximum IT = 100 ms, scan range 150 to 2000 *m/z*; dd-MS2 settings: resolution = 17,500, AGC target = 1x10^5^, maximum IT = 50 ms, loop count = 5, isolation window 1.5 *m/z*, isolation offset 0.5 *m/z*, stepped Normalized Collision Energy (NCE) = 20, 40, 60; dd settings: minimum AGC target = 8x10^3^, exclude isotopes ON, dynamic exclusion = 1 s.

### LC-MS analysis

For *S. peucetius* samples, targeted absolute quantification of daunorubicin production was performed using a dilution series of daunorubicin standards run alongside the culture extracts. A calibration curve was generated from these standards to calculate daunorubicin concentrations using the Quantitative Browser in Thermo Xcalibur (v4.3.73.11, Thermo Fisher Scientific). To account for variations in biomass, calculated concentrations were normalized against total DNA concentration, quantified as described above.

For untargeted metabolomics, data processing was performed using Compound Discoverer (v3.3.3.200, Thermo Fisher Scientific). The processing pipeline included MS spectra alignment, compound detection and grouping, gap filling, background compound filtering (using culture medium blanks), elemental composition prediction, and differential production analysis. Principal component analysis (PCA), volcano plots, and peucemycin peak areas for relative quantification were extracted from the Compound Discoverer output and replotted using R.

Untargeted metabolomics analysis of *S. coelicolor* samples was performed in the same manner, with the addition of a targeted screen for known *S. coelicolor* metabolites using a mass list manually compiled from the literature (**Supplementary Data 8**). Compounds with a calculated accurate mass within 5 ppm of the compounds in the mass list were labelled in the dataset for further analysis. Of these, metabolites with available MS/MS data were investigated to validate their identity where possible through manual MS/MS data comparison against the literature or GNPS networking. Peak areas of selected metabolites within that group were extracted for relative quantification. To account for variations in biomass, peak areas for these known metabolites were normalized against the wet biomass weight of each sample. The output of the Compound Discoverer metabolomic analyses of *S. coelicolor* and *S. peucetius* can be found in **Supplementary data 7 and 9**, respectively.

Mass spectral molecular networking was performed for the metabolomes of both *S. coelicolor* and *S. peucetius* strains using positive-mode MS/MS data. Raw data files (.raw) were converted to the open .mzML format and uploaded to the Global Natural Products Social Molecular Networking (GNPS) platform (http://gnps.ucsd.edu) for processing ^68^.

The molecular networking workflow was executed using the standard online pipeline. MS/MS spectra were first pre-filtered by removing all fragment ions within ± 17 Da of the precursor *m/z*. The spectra were then window-filtered to retain only the top six fragment ions within a ± 50 Da window across the spectrum. Precursor and MS/MS fragment ion mass tolerances were both set to 0.02 Da. To eliminate media artefacts, uninoculated culture media blanks were included in the analysis, and any matching spectra found in these control samples were filtered out prior to network generation.

Network edges were filtered to require a cosine score greater than 0.7 and at least 6 matched peaks. Furthermore, edges between two nodes were retained only if each node appeared in the other’s top 10 most similar nodes. The maximum allowable size for a molecular family was set to 100 nodes; for larger networks, the lowest-scoring edges were iteratively removed until the family size fell below this threshold.

Spectral libraries within GNPS were searched using input data filtered in the same manner, requiring a minimum library match cosine score of 0.7 and at least 6 matched peaks. The resulting networks were exported as a .graphml file and visualized using Cytoscape (v3.10.4) ^69^.

### RNA Extraction and Sequencing

*Streptomyces* cultures (30 mL, YT) were inoculated with ∼10^6^ spores in sprung flasks and incubated at 28 °C (250 rpm). Cells were harvested at 2 and 5 days, washed with 1x PBS, and snap-frozen at -70 °C. Mycelia were lysed in a cocktail of acid-phenol (pH 4.3), chloroform:isoamyl alcohol (24:1), and Qiagen RLT buffer using Lysing Matrix B (MP Biomedicals) and a FastPrep-24 homogenizer (two cycles of 30 s at 6.0 m/s with intermediate cooling on ice). After centrifugation (16,000 x *g*, 15 min, 4 °C), RNA was purified from the supernatant using a RNeasy Mini Kit (Qiagen) with on-column DNase I digestion, followed by a secondary genomic DNA removal step via a TURBO DNA-free Kit (Invitrogen). High-throughput library preparation and sequencing were performed by Novogene. RNA-seq data were analyzed as described below.

### RNA-seq Analysis

Data analysis was conducted as described previously ^70^. Read quality was assessed using FastQC, and transcripts were aligned to the *S. coelicolor* M145 genome (AL645882.2) using Bowtie2. The resulting alignments were sorted and indexed using Samtools. A custom Perl script was used to generate a SAF file for the *S. coelicolor* genome and read counts per gene were quantified using the featureCounts tool within the Rsubread package. Differential expression analysis was performed using edgeR in R. Count data were modeled using a quasi-likelihood negative binomial generalized log-linear fit via the glmQLFit function, and gene-wise statistical significance was evaluated using glmQLFTest. Differentially expressed genes (DEGs) were defined using a threshold of |log2 fold-change| ≥ 1 and a false discovery rate (FDR) adjusted *P*-value of ≤ 0.05 (**Supplementary Data 2**).

### Functional Annotation and Enrichment Analysis

Genes belonging to the actinorhodin (BGC0000194.5), undecylprodigiosin (BGC0001063.5) and coelimycin P1 (BGC0000038.5) BGCs were identified using the MIBiG database v4.0 ^71^.

Clusters of Orthologous Groups (COGs) functional categories were assigned to all proteins encoded by quantified transcripts using eggNOG-mapper against version 5.0.2 of the eggnog database ^72,73^ via the Galaxy web platform (Version 2.1.13+galaxy0) (The Galaxy Community, 2024). Transcripts lacking an identifiable seed ortholog or an assigned COG term were classified as "No ortholog found" or "No annotation," respectively, while genes with multiple COG assignments were counted toward each applicable category. Overrepresentation of functional classes among significantly upregulated or downregulated transcripts was assessed using an enrichment factor, defined as the ratio of the percentage of a given COG term in differentially expressed transcripts to its percentage in the total dataset.

#### COG class definition

**C**, energy production and conversion; **E**, amino acid transport and metabolism; **F**, nucleotide transport and metabolism; **G**, carbohydrate transport and metabolism; **H**, coenzyme transport and metabolism; **J**, translation, ribosomal structure and biogenesis; **K**, transcription; **L**, replication, recombination and repair; **M**, cell wall/membrane/envelope biogenesis; **O**, posttranslational modification, protein turnover, and chaperones; **P**, inorganic ion transport and metabolism; **Q**, secondary metabolites biosynthesis, transport and catabolism; **S**, function unknown; **T**, signal transduction mechanisms; **U**, intracellular trafficking, secretion, and vesicular transport; **V**, defense mechanisms. “No annotation” indicates proteins without an assigned COG function, and “No orthologue” indicates proteins without an identifiable orthologue in the reference genome.

### Chromatin co-immunoprecipitation with deep sequencing (ChIP-seq)

ChIP-seq was performed on biological duplicates as described previously ^74^ using *S. coelicolor* Δ*afsR* strains complemented *in trans* with wild-type or autoactive AfsR^GV^-1xFLAG under its native promoter at the ΦBT1 integration site (MJ187/MJ213). 30 ml YT cultures were inoculated with ∼10^6^ spores and grown at 28 °C (250 rpm) for 2 or 5 days.

Formaldehyde was added to a final concentration of 1% (v/v), and the cultures were incubated for 30 min to induce cross-linking, after which the reaction was quenched with 125 mM glycine for 5 min. Harvested mycelia were washed twice with 1x PBS and snap-frozen at -70 °C.

Frozen pellets were resuspended in lysis buffer (20 mM KHEPES pH 7.9, 50 mM KCl, 10% v/v glycerol) supplemented with 15 mg/mL lysozyme and Roche cOmplete protease inhibitor, and incubated at 37 °C for 25 min. Following dilution with an equal volume of lysis buffer, the samples were sonicated (13 cycles of 15 s on/off at 8 µm) on ice to yield a DNA shear size of ∼500 bp, as verified via agarose gel electrophoresis. Lysates were cleared by centrifugation (16,000 x *g*, 15 min, 4 °C), and the supernatant was adjusted to immunoprecipitation conditions by adding 10 µL of 1 M Tris (pH 8.0), 20 µL of 5 M NaCl, and 10 µL of 10% NP40 per milliliter.

Immunoprecipitation was performed overnight at 4 °C using anti-FLAG M2 affinity gel beads (Sigma) pre-equilibrated in IPP150 buffer (10 mM Tris-HCl pH 8.0, 150 mM NaCl, 0.1% NP40). Beads were washed five times with IPP150 buffer and twice with TE buffer. Bound complexes were sequentially eluted at 65 °C using elution buffer (50 mM Tris-HCl pH 8.0, 10 mM EDTA, 1% SDS) and TE containing 1% SDS. The pooled eluates were incubated overnight at 65 °C to reverse cross-links, and the DNA was purified using a QIAquick PCR Purification Kit (Qiagen). DNA concentration was quantified using a Qubit fluorometer, and samples were submitted to Azenta for library construction and sequencing.

### ChIP-seq Analysis

For analysis of the ChIP-seq data, a reference genome of *S. coelicolor* A3(2) SCP1^-^ SCP2^-^ Δ*afsR* ϕBT1::*afsR* was first constructed by introducing the Δ*afsR* deletion according to Floriano and Bibb ^75^ into the *S. coelicolor* M145 genome (AL645882.2) and inserting the sequence of the plasmid pMJ23, at the ϕBT1attachment site (SCO4848).

Sequencing reads were aligned to *S. coelicolor* A3(2) SCP1^-^ SCP2^-^ Δ*afsR* ϕBT1::*afsR* using the subread-align command of the Rsubread package ^76^ and Samtools v1.22 was used to sort, index, and compute nucleotide-resolution read depth from the resulting BAM files. Local enrichment was calculated across 30-nucleotide sliding windows, moving in 15-nucleotide steps, by comparing read density in the window to the density of reads in 3000 nucleotides surrounding the window. Control-subtracted local enrichment was computed by subtracting the local enrichment in M513 controls from the local enrichment in experimental samples. Regions in which ChIP local enrichment was ≥2 and control-subtracted local enrichment was positive were recorded. These enrichment regions were associated with the nearest genes on either side. For any given gene, the site of highest associated enrichment was considered the binding site for the gene.

To identify significantly enriched regions compared to the M513 control, the following cutoffs were applied: adjusted P-value (computed using R functions pnorm and p.adjust) of the control-subtracted local enrichment ≤ 0.05 and log_2_ fold change when comparing local enrichment in AfsR-FLAG/AfsR^GV^-FLAG to M513 controls > 1 at the site of highest associated enrichment. The list of identified enrichment sites is provided in Supplementary Dataset 3.

### Protein Expression and Purification

*E. coli* codon-optimized DNA fragments encoding wild-type *S. coelicolor* AfsR and its autoactive variant (AfsR^GV^) were synthesized (IDT) and cloned into pTB146 downstream of an N-terminal 6xHis-SUMO tag. Recombinant proteins were produced in *E. coli* BL21(DE3) grown in LB containing carbenicillin and chloramphenicol at 37 °C to an OD_600_ of 0.5-0.6. Expression of wild-type *afsR* was induced with 0.5 mM isopropyl-β-D-thiogalactopyranoside (IPTG) for 4 h at 30 °C, whereas expression of the autoactive variant *afsR^GV^* was induced with 0.4 mM IPTG overnight at 16 °C. Harvested cells were resuspended in Buffer A (50 mM Tris-HCl pH 8.0, 10% sucrose, 300 mM KI, 1 mM ATP, 10 mM imidazole, 1 mM TCEP) supplemented with Roche cOmplete protease inhibitors and Pierce universal nuclease, and lysed via sonication.

Lysates were cleared by centrifugation (25,312 x *g*, 40 min), and supernatants were loaded onto a 5 mL HisTrap Excel column (Cytiva). Bound proteins were eluted using a 10-500 mM imidazole linear gradient with Buffer B (Buffer A containing 500 mM imidazole). Target fractions were pooled based on SDS-PAGE analysis. For tag removal, pooled proteins were incubated with 6xHis-Ulp1 protease (100:1 molar ratio) during overnight dialysis at 4 °C against a dialysis buffer (50 mM Tris-HCl pH 8.0, 10% sucrose, 300 mM KI, 1 mM ATP, 1 mM DTT).

The cleaved His6-SUMO tag and His6-Ulp1 were removed by gravity-flow chromatography using HIS-select nickel affinity gel (Sigma-Aldrich). The untagged flow-through fraction containing AfsR and AfsR^GV^, respectively, was concentrated and subjected to size-exclusion chromatography using a HiLoad 16/600 Superdex 200 pg column (Cytiva) equilibrated in gel filtration buffer (Buffer A without imidazole). Eluted fractions were assessed for purity by SDS–PAGE before desired fractions were pooled, concentrated, and flash-frozen at -70 °C.

### AfsR Antibody Generation

To produce a polyclonal antibody against AfsR from *S. coelicolor*, recombinant, untagged AfsR was produced as described above and sent to Biosynth Laboratories Ltd (UK).

### Mass photometry

Mass photometry measurements of purified AfsR and AfsR^GV^ were performed using a Refeyn TwoMP mass photometer. Protein samples were diluted to 100 nM in assay buffer (50 mM Tris-HCl, pH 8.0, 300 mM KI, 1 mM ATP, 1 mM TCEP; filtered through a 0.22 μm membrane). To set the focal plane, 10 μL of assay buffer was loaded into the sample well and autofocus was engaged. Subsequently, 2–5 μL of 100 nM AfsR or AfsRGV, respectively, was added to the droplet and mixed to initiate data acquisition. Contrast interference movies were recorded for 60 s at room temperature using default acquisition settings in AcquireMP software (v2024 R2.1). All measurements were performed in triplicate. Contrast values were converted to molecular mass using a standard calibration curve, and mass distribution profiles were processed and analyzed using DiscoverMP (v2024 R2.1).

### Biolayer Interferometry

Protein-DNA binding kinetics were evaluated using biolayer interferometry on an Octet-N1 (Sartorius) instrument with High Precision Streptavidin biosensors (Octet SAX2 biosensors, Sartorius) pre-equilibrated in assay buffer (50 mM Tris-HCl pH 7.5, 200 mM NaCl, 1 mM MgCl_2_, 0.1 mg/mL BSA, 1 mM DTT, 0.05% Tween 20).

Double-stranded DNA targets were generated by annealing 5’-biotinylated oligonucleotides (IDT, **Supplementary Table S3**) at 98 °C for 5 min followed by slow cooling. Immobilization of 100 nM dsDNA onto the biosensors was performed using a baseline-association-dissociation cycle (30 s/120 s/120 s). For affinity measurements, sensors were incubated with a 1:1 serial titration of AfsR or AfsR^GV^ (40 nM to 5.12 μM), using a 30 s baseline, 300 s association, and 300 s dissociation profile in triplicate. Data were baseline-corrected using Octet N1 1.4 software, and steady-state binding signals extracted at 326 s were plotted against protein concentration. Equilibrium dissociation constants (K_D_) were determined in GraphPad Prism (Vers. 5) using a non-linear regression, one-site binding model.

### Statistical analysis

Statistical analyses and data visualization were performed in R v4.4.1 and RStudio 2024.09.0+375 using the packages dplyr (v1.2.1; Wickham et al., 2023), Rmisc (v1.5.1; Hope, 2022), ggplot2 (v4.0.4; Wickham, 2016) and GraphPad Prism 11.

For *S. coelicolor* metabolite data, biomass-normalized peak intensities were compared between strains MJ181 and MJ171. Normality and homogeneity of variances were evaluated using the Shapiro-Wilk test and F-test, respectively; statistical significance was determined using a Welch Two-sample t-test for normally distributed data or a Wilcoxon Rank-Sum test for non-parametric data.

For the *S. peucetius* metabolomics data, statistical analysis was performed on the production of peucemycin, hydroxy-peucemycin, and daunorubicin at each time point. The Levene test ^77^ and the Shapiro-Wilk test were used to check the homogeneity of variances and normality assumptions of the ANOVA model. When the data failed to meet these assumptions, a Kruskal-Wallis test was applied, followed by the Dunn test with a Benjamini-Hochberg correction to determine differences between individual strains. When the assumptions were met, a one-way ANOVA was applied, followed by Tukey’s honestly significant difference (HSD) post hoc test for pairwise comparisons.

### Declaration of Generative AI and AI-Assisted Technologies in the Writing Process

ChatGPT (OpenAI) and Gemini (Google) were used during manuscript preparation to improve grammar and readability and to assist with writing and refining code for bioinformatic analyses. The authors critically reviewed and edited all AI-generated suggestions and remain fully responsible for the manuscript content.

### Data availability

The ChIP-seq and RNA-seq data generated in this work have been deposited in the ArrayExpress database under the accession codes E-MTAB-17467 and E-MTAB-17362, respectively. GNPS outputs obtained as part of the metabolomics analysis can be viewed as follows: *S. coelicolor* https://gnps.ucsd.edu/ProteoSAFe/status.jsp?task=f7ebef248f3e46859bf8942d4e6cacb8, *S. peucetius* four-day samples https://gnps.ucsd.edu/ProteoSAFe/status.jsp?task=f5969eeedc9c46b2b8b24aa2e2d44e9c, and *S. peucetius* 6-day samples https://gnps.ucsd.edu/ProteoSAFe/status.jsp?task=f1671c3589c14094add1b663dd1c2554. LC-MS/MS files in the mzML format have been deposited in the MassIVE repository under accession number MSV000102706 (*S. coelicolor* MS files) and MSV000102712 (*S. peucetius* MS files).

Scripts used to perform the reciprocal BLAST analysis contained in this study are available on GitHub (https://github.com/ml-jordan/afsr-reciprocal-blast). Other data are included in the article and/or supplementary information.

## Supporting information

Combined Datasets S1-S6

## Acknowledgments

We thank Sophien Kamoun and Mark Buttner for helpful discussions and Matthew Herdman for preliminary protein purifications.

## Author contributions

Conceptualization: MLJ, SS

Methodology: MLJ, HW, NMV, KJS, AM, GC, MJB

Investigation: MLJ, HW, NMV, SS

Visualization: MLJ, HW, NMV, SS

Funding acquisition: SS

Project administration: SS

Supervision: AT, SS

Writing – original draft: MLJ, SS

Writing – review & editing: MLJ, HW, NMV, AT, SS

