## Supplementary Information for "A conserved NLR activation switch governs global transcriptional control of antibiotic biosynthesis in *Streptomyces*"

**This PDF file includes:**

Figures S1 to S10
Tables S1 to S3
SI References

**Other supporting materials for this manuscript include the following:**

Datasets S1 to S9

### Figures

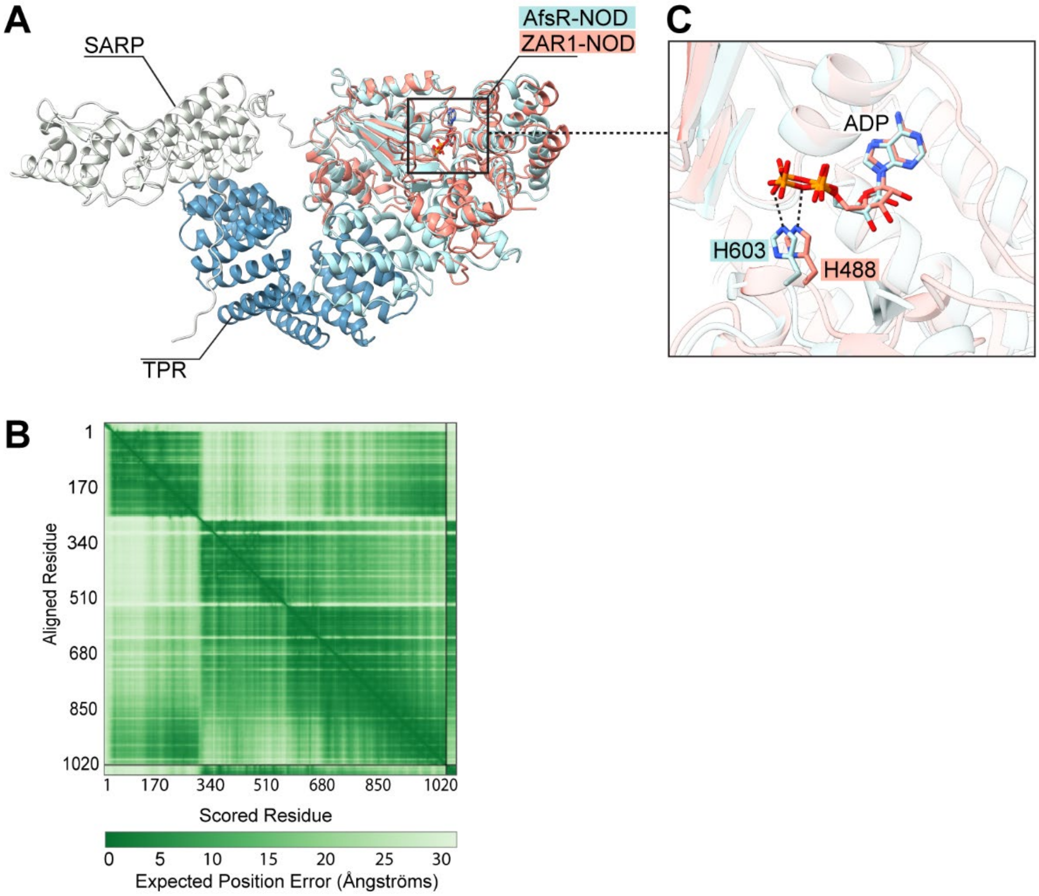


**Fig. S1. Structural conservation of autoactivity-associated residues in AfsR. (A)** Structural model of *S. coelicolor* AfsR with ADP generated with AlphaFold3. The three AfsR domains are shown: SARP effector domain (white), NOD module (light blue) and TPR, sensor domain (dark blue). The experimentally determined NOD fragment of *A. thaliana* ZAR1 (salmon, PDB: 6J5W) is superimposed on the NOD module of AfsR. **(B)** Predicted alignment error (PAE) plot for the AfsR-ADP model shown in (A). **(C)** Magnified view of the predicted ADP-binding pocket of AfsR, superimposed onto the ADP-binding pocket in the cryoEM structure of ZAR1 (PDB: 6J5W). The conserved histidine residues in AfsR (H603, light blue) and ZAR1 (H488, salmon) involved in nucleotide binding are shown as sticks and labeled. ADP molecules are shown as sticks.

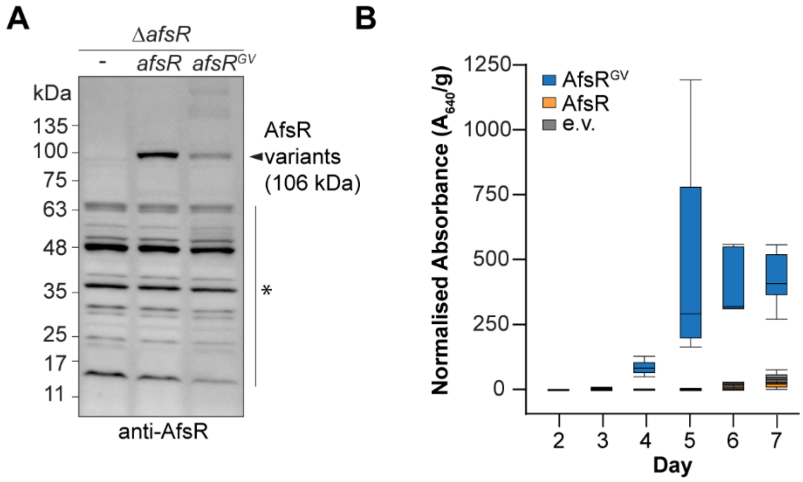


**Fig. S2. Autoactivation of AfsR increases actinorhodin production in *S. coelicolor*. (A)** Immunoblot analysis using an anti-AfsR antibody to detect the production of AfsR and AfsR$^{GV}$ in $\Delta$*afsR S. coelicolor* cells grown for two days in YT broth. Asterisk denotes non-specific bands. Shown is a representative image (n=3). **(B)** Quantification of actinorhodin production of $\Delta$*afsR S. coelicolor* cells carrying the empty vector (e.v., grey) or producing AfsR (orange) and AfsR$^{GV}$ (blue). Cells were grown in YT broth, and actinorhodin production was analyzed at the indicated time points. Production levels in the different strains were normalized by cell pellet weight. Limits of the boxplots represent the 25th–75th percentile, the horizontal line denotes the median, and the whiskers represent the minimum/maximum values (n = 6).

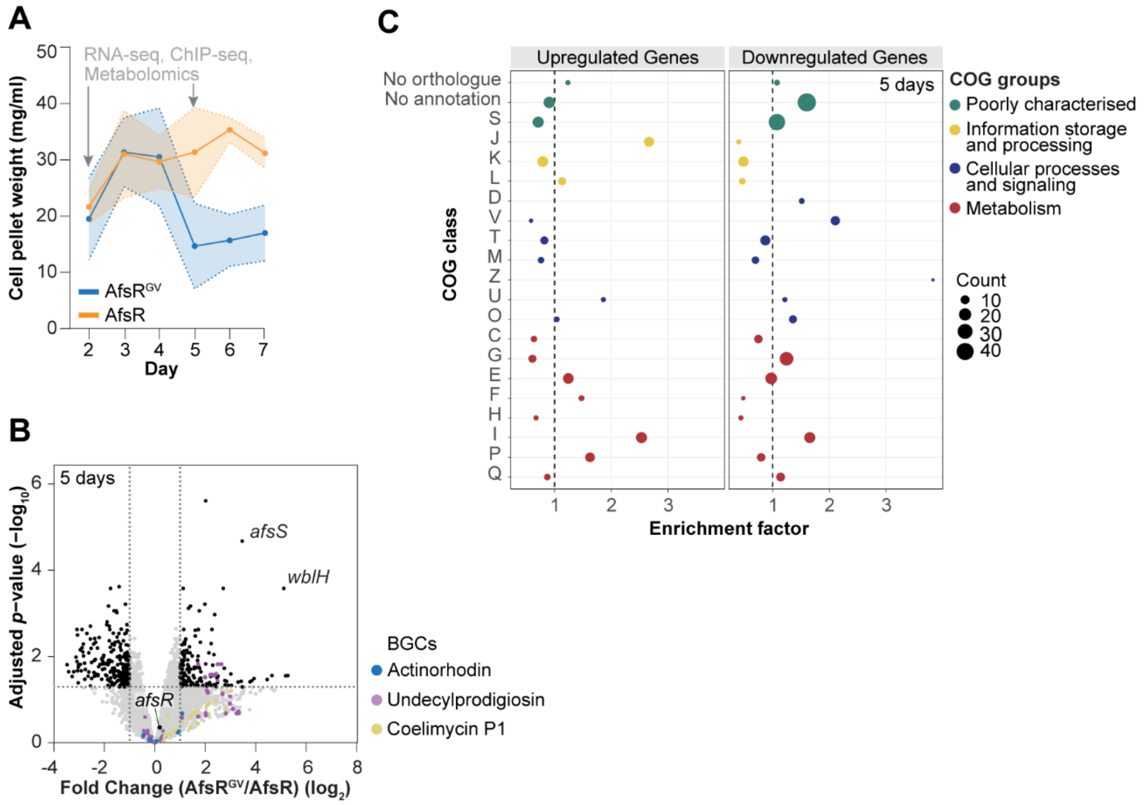


**Fig. S3. Expression of *afsR*$^{GV}$ induces upregulation of metabolism-associated genes.** **(A)** Growth curve of Δ*afsR S. coelicolor* complemented with either *afsR* (orange) or *afsR*$^{GV}$ (blue). Shown is the mean pellet weight, shaded area represents standard error (n=6). Arrows point at time points (2 and 5 days) used for RNA-seq, ChIP-seq and metabolomic analyses. **(B)** Volcano plot of differentially expressed genes (DEGs) at 5 days of cultivation in complemented *S. coelicolor* Δ*afsR/afsR*$^{GV}$ cells compared to Δ*afsR/afsR*. Data derived from 3 biological replicates. Dashed lines indicate significance thresholds. Selected genes and gene groups are identified by labels. **(C)** Analysis of Clusters of Orthologous Groups of proteins (COG) enriched in cells expressing *afsR*$^{GV}$ compared to *afsR* at 5 days. Bubble size represents the number of DEGs assigned to each COG class. See also **Fig. 2A-B** for results from 2-day samples. See Methods section for COG class definition.

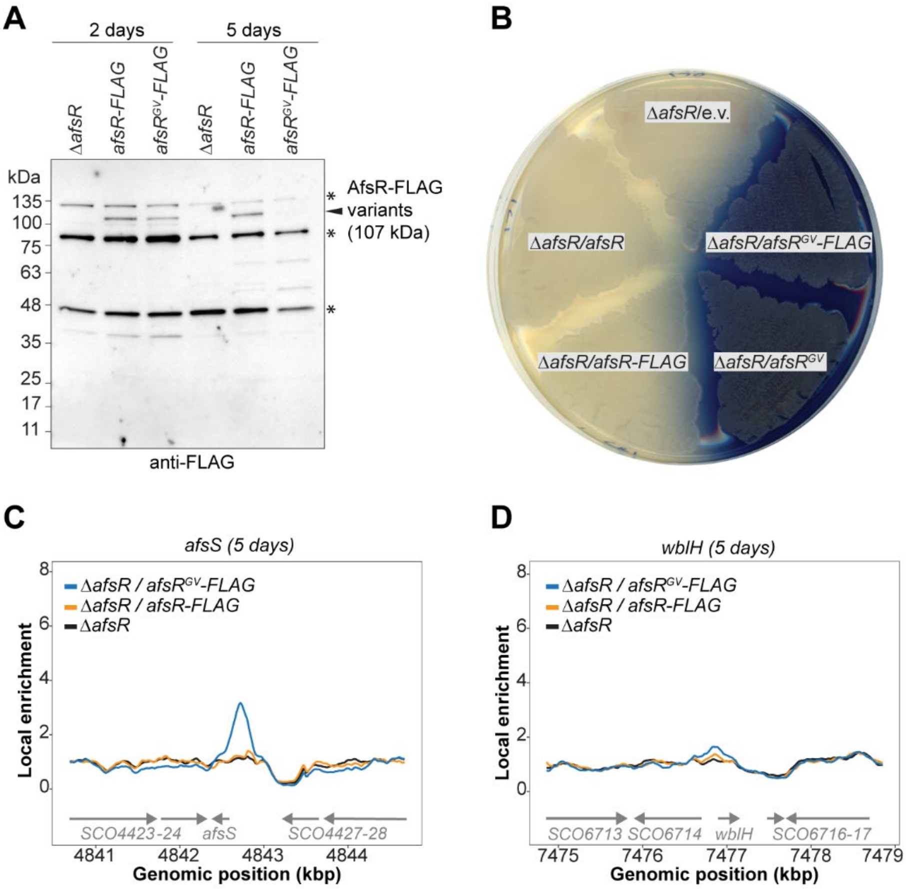


**Fig. S4: Characterization of FLAG-tagged AfsR variants. (A)** Immunoblot analysis using an anti-FLAG antibody to detect the production of AfsR-FLAG and AfsR$^{GV}$-FLAG variants in Δ*afsR S. coelicolor* cells grown for 2 and 5 days in YT. Asterisk denotes non-specific bands. Shown is a representative image (n=3). **(B)** Actinorhodin production (blue) of *S. coelicolor* Δ*afsR* cells complemented with *afsR*, *afsR$^{GV}$*, *afsR*-FLAG or *afsR$^{GV}$*-FLAG expressed from the native promoter, or the empty vector (e.v.). Representative image of strains grown on Difco nutrient agar and imaged after 3 days of incubation. **(C)** and **(D)** ChIP-seq profiles showing AfsR$^{GV}$-FLAG and AfsR-FLAG enrichment at the *S. coelicolor afsS* (C) and *wblH* (D) promoter region at 5 days. See **Fig. 2C-D** for results from 2-day samples. Analysis was performed on 2 biological replicates.

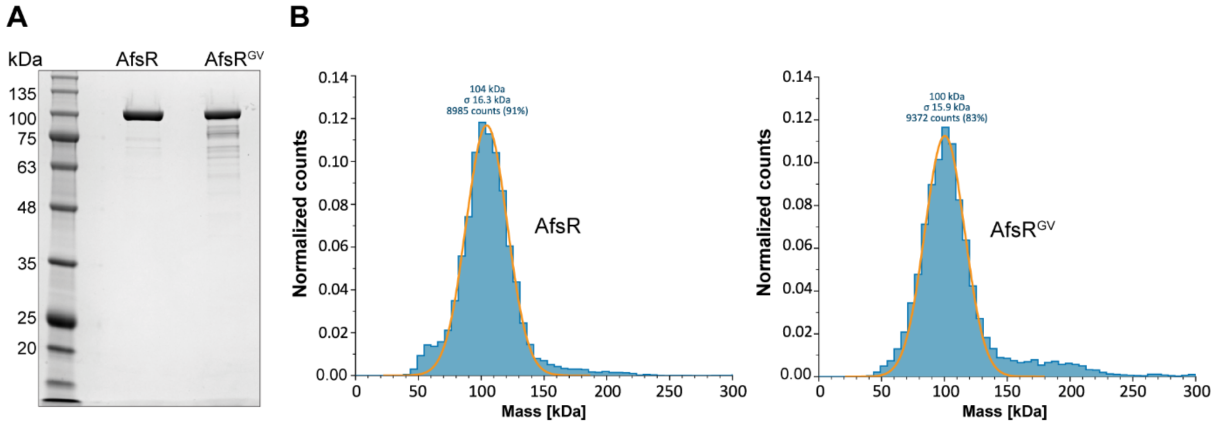


**Fig. S5. Purification of AfsR and AfsR$^{GV}$. (A)** Coomassie-stained SDS gel with purified AfsR and AfsR$^{GV}$ (106 kDa). **(B)** Mass photometry analysis of purified AfsR and AfsR$^{GV}$, indicating that both proteins are monomeric in solution. The molecular mass of the particles, corresponding standard deviation (σ) and the number of particles counted are indicated. Shown are representative results of triplicate experiments.

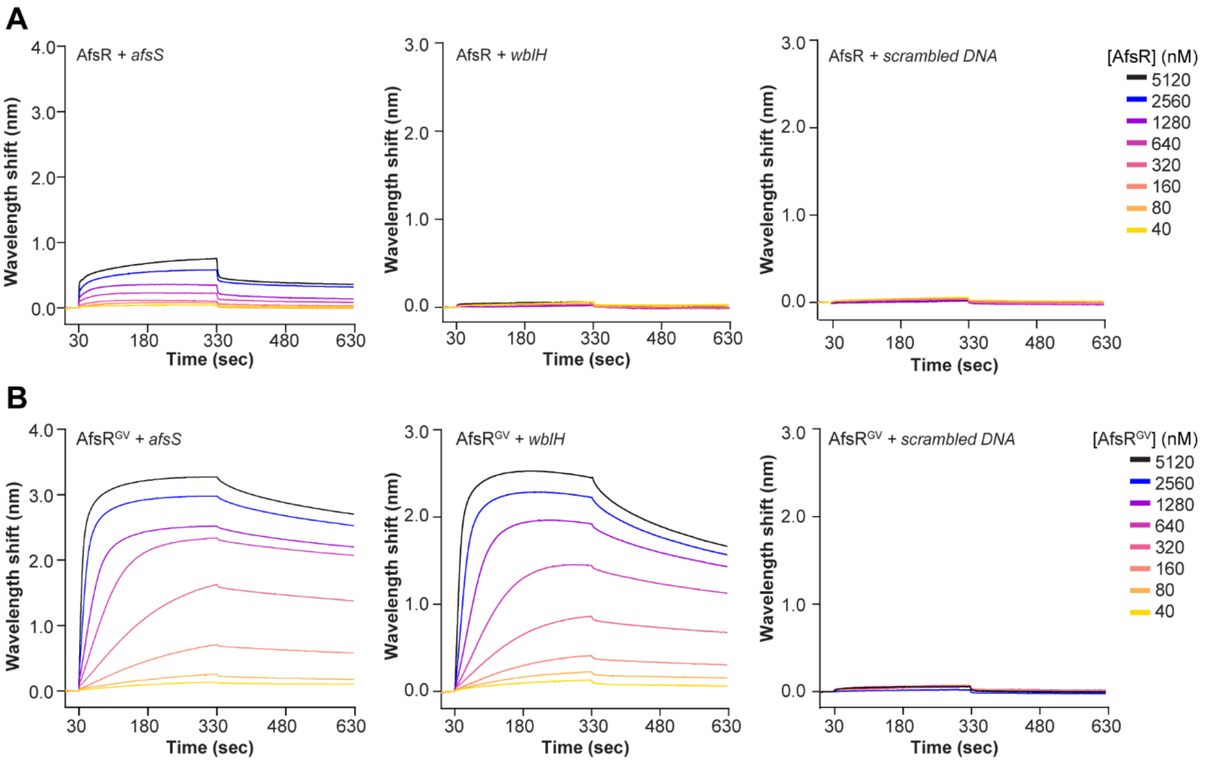


**Fig. S6. Biolayer interferometry analysis of DNA-binding behavior of (A)** AfsR and **(B)** AfsR$^{GV}$. A double-stranded, biotinylated DNA fragment (42 bp) containing the binding motif in the *afsS* and *wblH* promoter region or a randomized (scrambled DNA) sequence was immobilized on a streptavidin-coated biosensor and probed with the indicated concentrations of AfsR or AfsR$^{GV}$. The graphs show the mean results of 3 independent experiments.

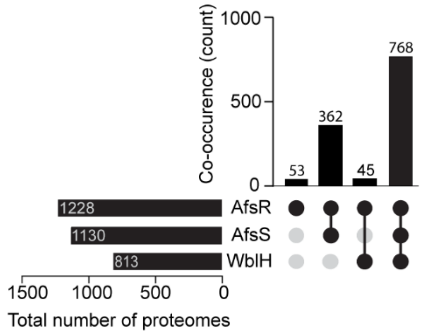


**Fig. S7**. **Distribution of WblH and AfsS in *Streptomyces*.** Upset plot shows the co-occurrence of WblH and AfsS in *Streptomyces* proteomes (n=1228) that contain AfsR.

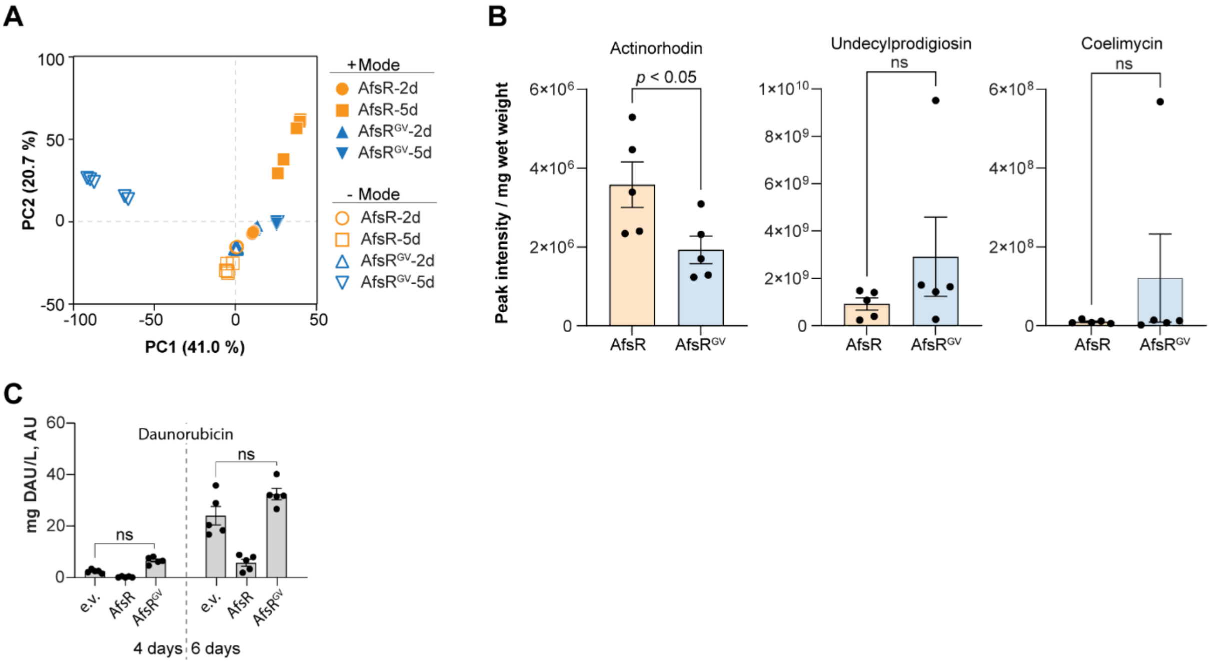


**Fig. S8. Metabolomic response to autoactive AfsR$^{GV}$ in *S. coelicolor* and *S. peucetius*.** **(A)** Principal Component Analysis (PCA) of the metabolome of *S. coelicolor* Δ*afsR* complemented with *afsR* (orange) or *afsR$^{GV}$* (blue). Strains were grown in YT broth; cultures were sampled after 2 (2d) and 5 days (5d) of fermentation and analyzed by LC-MS using Compound Discoverer 3.3 (Thermo). Data were obtained from 5 biological replicates. **(B)** Bar charts comparing the production of selected metabolites in *S. coelicolor* Δ*afsR* expressing *afsR* or *afsR$^{GV}$* at 2 days of fermentation. The MS peak area was normalized by the cell pellet weight for each sample. Error bars represent the standard error (n=5). Statistical significance ($p$ value) was determined using the Wilcoxon rank sum test. See also **Fig. 3A-C**. **(C)** Bar charts comparing the production of daunorubicin detected in culture supernatants of *S. peucetius* expressing *afsR or afsR$^{GV}$* from the native promoter. e.v. is an empty vector control. MS peak areas are normalized by culture growth (colorimetric DNA measurement). Data are derived from 4 and 6 days of fermentation in SFM broth, with 5 biological replicates per strain. Error bars represent the standard error. Statistical significance (adjusted $p$ value) was determined using Kruskal-Wallis tests followed by a Dunn test with BH correction (4-day samples) and one-way ANOVA followed by a Turkey's HSD post-hoc test (6-day samples); ns, not significant.

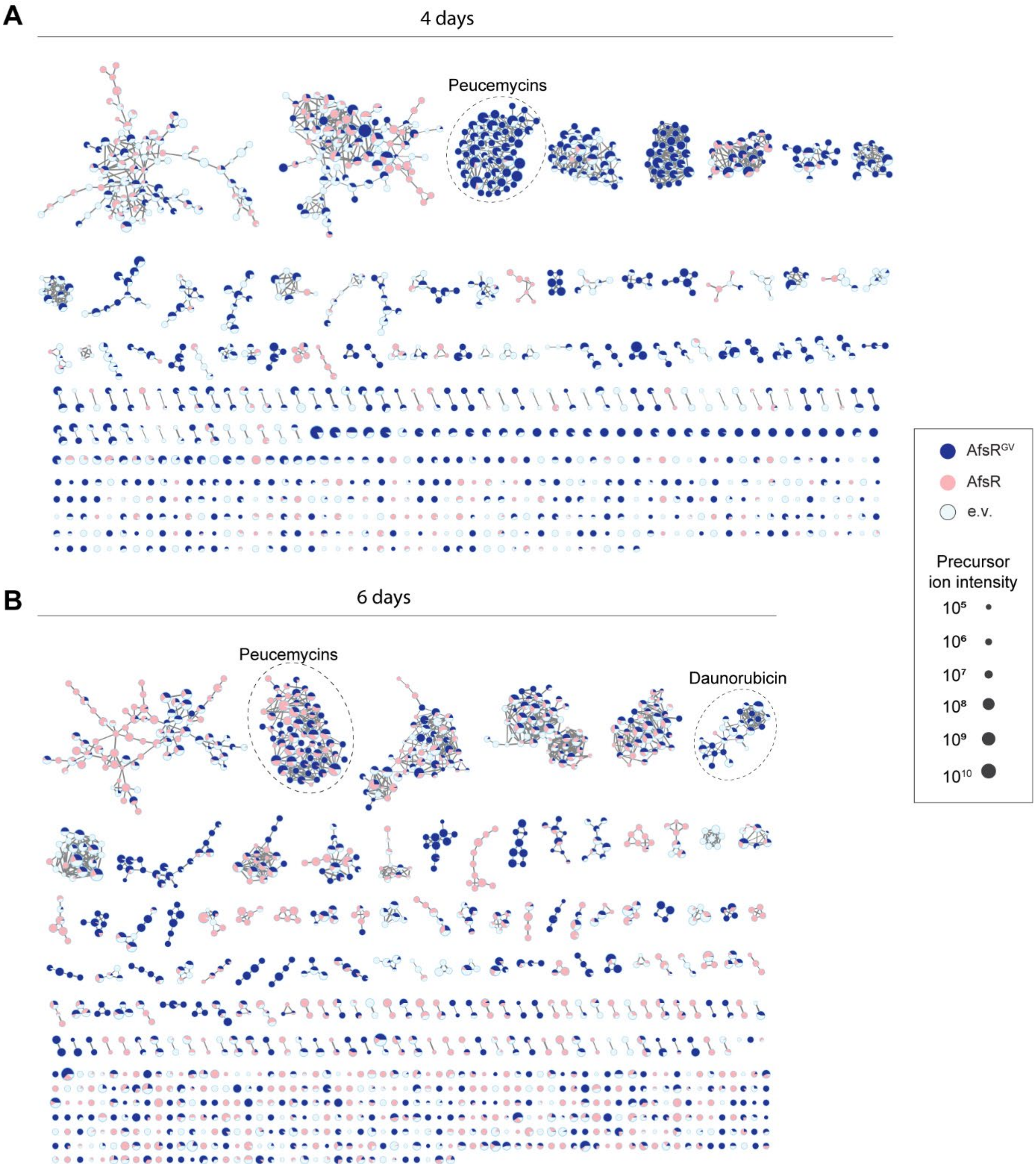


**Fig. S9. Molecular structural diversity of molecules detected in culture supernatants of *S. peucetius* expressing *afsR* variants under the control of the native promoter**. Mass spectral molecular network generated from positive MS data acquired after 4 and 6 days of fermentation in SFM broth. Nodes represent metabolites, with sizes proportional to their precursor ion intensity, and pie charts indicate relative abundance in cells expressing *afsR* and *afsR*$^{GV}$ or in the empty-vector control (e.v.). Edges are weighted by MS/MS cosine similarity scores. Dashed circles indicate groups of known molecules. MS Peak areas are normalized by culture growth (colorimetric DNA measurement). Data are derived from 5 biological replicates per strain.

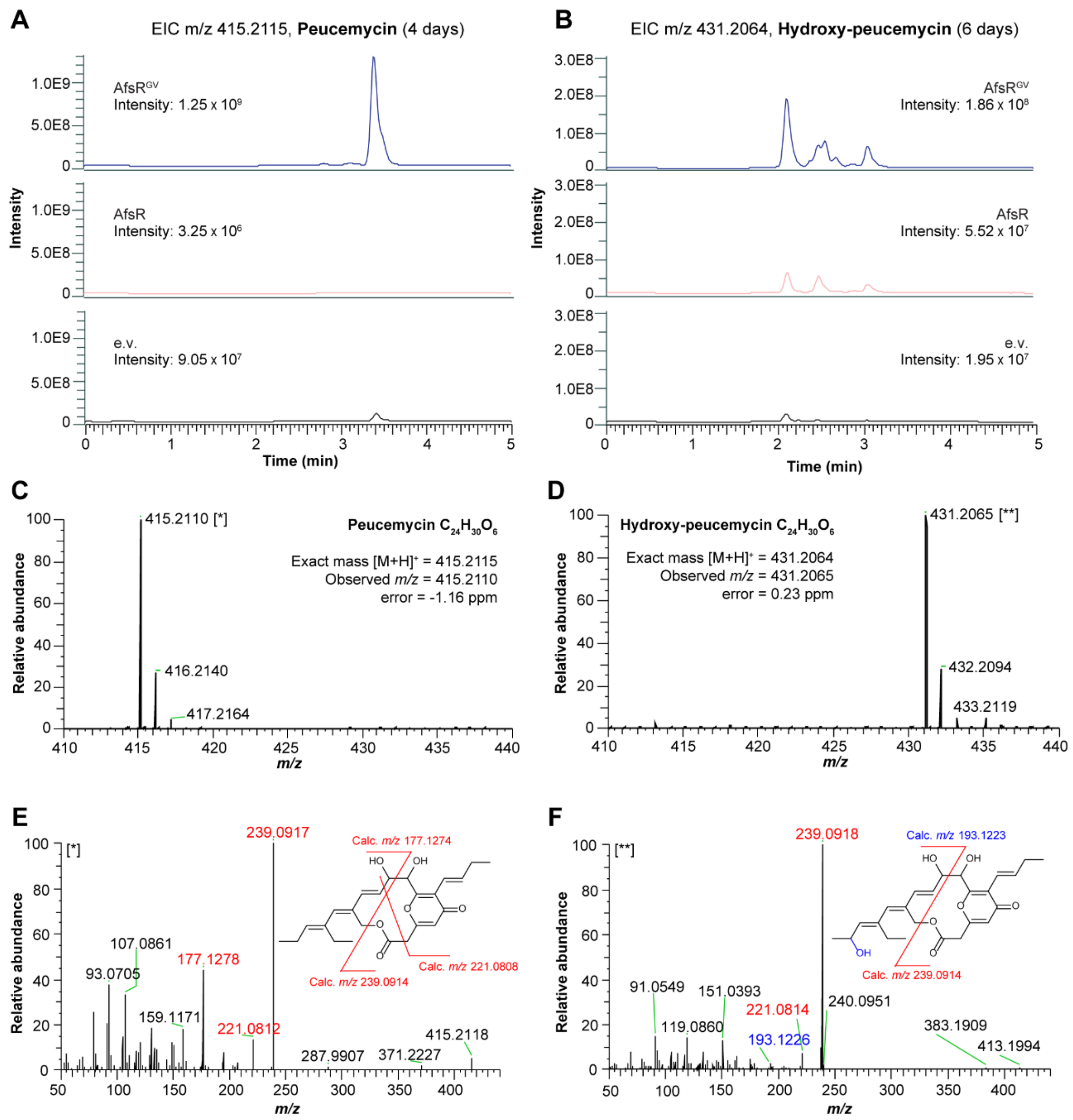


**Fig. S10. Validation of peucemycin production in culture supernatant extracts of *S. peucetius* expressing *afsR* variants under the control of its native promoter.** Extracted ion chromatograms (EICs) for **(A)** Peucemycin *m/z* 415.2115 at 4 days and **(B)** Hydroxy-peucemycin *m/z* 431.2064 at 6 days of fermentation in SFM broth; e.v. is the empty vector control. Chromatograms are scaled to the maximum observed intensity for each compound, and the peak intensity per sample is listed next to its corresponding chromatogram. Data are derived from 5 biological replicates per strain. **(C-D)** MS spectra for (C) peucemycin and (D) hydroxy-peucemycin, showing the mass difference between their theoretical and detected masses. **(E)** and **(F)** fragmentation (MS/MS) spectra and proposed structures of (E) peucemycin [*] and (F) hydroxy-peucemycin [**]. Characteristic fragments are highlighted in red and blue in the MS/MS spectra and structures.

### Tables

**Table S1.** List of bacterial strains used in this work.

| Strain | Description | Use | Source |
|---|---|---|---|
| *S. coelicolor* | *S. coelicolor* M145, *SCP1*$^-$, *SCP2*$^-$ | Wild-type. | (1) |
| M513 | *S. coelicolor* M145 Δ*afsR* | Background strain, ChIP-seq analysis. | (2) |
| MJ170 | *S. coelicolor* M513 attB ΦBT1::pIJ10770 Hyg$^R$ | Metabolomics, RNA-seq analysis. | This study. |
| MJ171 | *S. coelicolor* M513 attB ΦBT1::*afsR* Hyg$^R$ | Metabolomics, RNA-seq analysis. | This study. |
| MJ181 | *S. coelicolor* M513 attB ΦBT1::*afsR*(H603G/D604V) Hyg$^R$ | Metabolomics, RNA-seq analysis. | This study. |
| MJ187 | *S. coelicolor* M513 attB ΦBT1::*afsR-flag* Hyg$^R$ | ChIP-seq analysis. | This study. |
| MJ213 | *S. coelicolor* M513 attB ΦBT1::*afsR*(H603G/D604V)-*flag* Hyg$^R$ | ChIP-seq analysis. | This study. |
| SS594 | *S. coelicolor* M513 attB ΦBT1::*afsR-3xflag* Hyg$^R$ | Phenotypic analysis. | This study. |
| SS598 | *S. coelicolor* M513 attB ΦBT1::*afsR*(H603G/D604V)-*3xflag* Hyg$^R$ | Phenotypic analysis. | This study. |
| *S. peucetius* | *S. peucetius* var. caesius ATCC 27952 | Wild-type. | DSMZ. (DSM 41231) |
| MJ300 | *S. peucetius* attB ϕC31::pIJ12552 Apra$^R$ | Metabolomics analysis. | This study. |
| MJ218 | *S. peucetius* attB ϕC31::Sp-*afsR* Apra$^R$ | Metabolomics analysis. | This study. |
| MJ220 | *S. peucetius* attB ϕC31::Sp-*afsR*(H594G/D595V) Apra$^R$ | Metabolomics analysis. | This study. |
| *E. coli* TOP10 | F$^-$ *mcrA* Δ(*mrr-hsdRMS-mcrBC*) ϕ80*lacZ*ΔM15 Δ*lacX74 recA1 araD*139 Δ(*ara-leu*)7697 *galU galK* λ- *rpsL*(Str$^R$) *endA1 nupG* | Standard cloning strain for generating plasmids used in this study. | Lab stock |
| *E. coli* Rosetta(DE3) | *F*$^-$ *ompT hsdS*$_B$(*r*$_B^-$ *m*$_B^-$) *gal dcm (DE3)* pRARE *(Cam*$^R$*)* | Host strain for protein over-expression plasmids. | Lab stock |
| ET12567/ pUZ8002 | *E. coli* F$^-$ *dam*-13::Tn9 *dcm*-6 *hsdM hsdR* recF143::Tn10 *galK2 galT22 ara*-14 *lacY*1 *xyl*-5 *leuB*6 *thi*-1 *tonA*31 *rpsL*136 *hisG*4 *tsx*-78 *mtl*-1 *glnV*44 Cam$^R$ Kan$^R$ | Host strain for introducing plasmids into *Streptomyces* spp. via conjugation. | (3) |

**Table S2.** List of plasmids used in this work.

| Vector | Description | Source |
|---|---|---|
| pIJ10770 | Cloning vector for expression of gene under the native promoter after conjugal transfer *from E. coli* to *Streptomyces* spp. Plasmid integrates specifically at the ΦBT1 phage integration site on the *S. coelicolor* chromosome. Hyg$^{R}$. | (4) |
| pIJ12552 | Cloning vector for expression of gene under the constitutive *ermE** promoter after conjugal transfer from *E. coli* to *Streptomyces* spp. Plasmid integrates specifically at the ΦC31 phage integration site on the *S. peucetius* chromosome. Apra$^{R}$. | Emma Sherwood, unpublished |
| pKF351 | Cloning vector for expression of gene under the native promoter after conjugal transfer from *E. coli* to *Streptomyces* spp. Plasmid integrates specifically at the ΦC31 phage integration site on the *S. peucetius* chromosome. Apra$^{R}$. | (4) |
| St6F11 | Cosmid vector containing coding sequence for *S. coelicolor* AfsR (SCO4426). | John Innes Centre http://strepdb.streptomyces.org.uk |
| pTB145 | Plasmid encoding the 6xHis-Ulp1 protease for expression and purification. Carb$^{R}$. | (5) |
| pTB146 | Plasmid encoding a 6xHis-SUMO tag for cloning genes for protein production and purification. Carb$^{R}$. | (5) |
| pMJ23 | *S. coelicolor* AfsR with its native promoter cloned into pIJ10770 for integration at the ΦBT1 site. Hyg$^{R}$. | This study. |
| pMJ35 | *S. coelicolor* AfsR containing the H603G/D604V mutations with its native promoter cloned into pIJ10770 for integration at the ΦBT1 site. Hyg$^{R}$. | This study. |
| pMJ52 | *S. coelicolor* AfsR with a C-terminal 1x FLAG tag with its native promoter cloned into pIJ10770 for integration at the ΦBT1 site. Hyg$^{R}$. | This study. |
| pMJ62 | *S. coelicolor* AfsR containing the H603G/D604V mutations with a C-terminal 1x FLAG tag with its native promoter cloned into pIJ10770 for integration at the ΦBT1 site. Hyg$^{R}$. | This study. |
| pMJ73 | *S. peucetius* AfsR with its native promoter cloned into pKF351 for integration at the ΦBT1 site. Apra$^{R}$. | This study. |
| pMJ75 | *S. peucetius* AfsR containing H594G/D595V with its native promoter cloned into pKF351 for integration at the ΦBT1 site. Apra$^{R}$. | This study. |
| pSS753 | *S. coelicolor* AfsR codon-optimised for *E. coli* expression cloned into pTB146. Carb$^{R}$. | This study. |
| pSS756 | *S. coelicolor* AfsR containing H603G/D604V mutations codon-optimised for E. coli expression cloned into pTB146. Carb$^{R}$. | This study. |

| | | |
|---|---|---|
| pSS766 | Cloning vector for expression of gene under the native or *ermE** promoter with a C-terminal 3xFLAG tag after conjugal transfer *from E. coli* to *Streptomyces* spp. Plasmid integrates specifically at the ΦBT1 phage integration site on the *S. coelicolor* chromosome. $Hyg^R$. | This study. |
| pSS772 | *S. coelicolor* AfsR with a C-terminal 3xFLAG tag with its native promoter cloned into pSS766 for integration at the ΦBT1 site. $Hyg^R$. | This study. |
| pSS778 | *S. coelicolor* AfsR containing the H603G/D604V with a C-terminal 3xFLAG tag with its native promoter cloned into pSS766 for integration at the ΦBT1 site. $Hyg^R$. | This study. |

**Table S3.** List of oligonucleotides used in this work.

| Name | Description | Sequence |
|---|---|---|
| MJ18 AfsR NP F | Amplification of *S. coelicolor afsR* along with its native promoter for insertion into pIJ10770. | CATCAGCAAAAGGGGATGATAAGTTTATCAAGCTTGCCTCCCCCGCCGCACA |
| MJ19 AfsR NP R | Amplification of *S. coelicolor afsR* along with its native promoter for insertion into pIJ10770. | ACCTAGGCTTAAGTCGCGAATCGATGATCATATGTCACCGCGCCACACTGCGAGC |
| MJ72 CA34 F1 R | Mutagenic primer for inserting H603G/D604V double mutation into *S. coelicolor* AfsR via Gibson assembly when cloning into pIJ10770. | AGGCGGACCAGGACGCCGAAGCGGTAGCG |
| MJ73 CA34 F2 F | Mutagenic primer for inserting H603G/D604V double mutation into *S. coelicolor* AfsR via Gibson assembly when cloning into pIJ10770. | GGCCGCTACCGCTTCGGCGTCCTGGTCCGCCTC |
| MJ150 AfsR R FLAG | Amplification of *S. coelicolor afsR* along with its native promoter for insertion into pIJ10770 with a C-terminal 1x FLAG tag. | ACCTAGGCTTAAGTCGCGAATCGATGATCATATGTCACTTATCGTCGTCATCCTTGTAATCCCGCGCCACACTGCGAGC |
| MJ195 AfsR_p NP GA F | Amplification of *S. peucetius afsR* along with its native promoter for insertion into pKF351. | GGACCGGATGAATTCACTTGGATCCTCATTCTAGAAGGTGAAGGAGATCAGCAAGGAGCTCG |
| MJ196 AfsR_p NP GA R | Amplification of *S. peucetius afsR* along with its native promoter for insertion into pKF351. | ACTAACGTCTGGAAAGACGACAAAACTTTAGATCTTCAGGCAGCCCTGAGCGGAG |
| MJ221 SpAfsR PM F | Mutagenic primer for amplifying *S. peucetius afsR* for insertion of the H594G/D595V mutations. | GGACCGGATGAATTCACTTGGATCCTCATTCTAGAAGGTGAAGGAGATCAGCAAGGAGCTCG |
| MJ222 SpAfsR PM R | Mutagenic primer for amplifying *S. peucetius afsR* for insertion of the H594G/D595V mutations. | ACTAACGTCTGGAAAGACGACAAAACTTTAGATCTTCAGGCAGCCCTGAGCGGAG |
| MJ231 SpAfsR PM34 F1 R | Mutagenic primer for insertion of the H594G/D595V mutations into *S. peucetius afsR*. | GAGACGCACAAGGACGCCGTACCGGTACCGGCCCGG |
| MJ232 SpAfsR PM34 F2 F | Mutagenic primer for insertion of the H594G/D595V mutations into *S. peucetius afsR*. | CGGTACCGGTACGGCGTCCTTGTGCGTCTCTACGCGCG |

| | | |
|---|---|---|
| SS1839 | Amplification of codon-optimised *S. coelicolor afsR* for insertion into pTB146 for protein expression and purification. | ATTGAGGCTCACAGAGAACAGATTGG TGGTGATGGTGGTCCACGCGTTCCAG AAC |
| SS1840 | Amplification of codon-optimised *S. coelicolor afsR* for insertion into pTB146. | TCGAGTGCGGCCGCAAGCTTGTCGAC GGAGTTATCGAGCTACTGATCTCGCTA AGAGTGC |
| SS1219 | Amplification of the pTB146 vector for insertion of codon-optimised *afsR* or *afsR*$^{GV}$ for protein expression and purification. | CTCCGTCGACAAGCTTGCGG |
| SS1220 | Amplification of the pTB146 vector for insertion of codon-optimised *afsR* or *afsR*$^{GV}$ for protein expression and purification. | ACCACCAATCTGTTCTCTGTGAGCC |
| SS1911 | Amplification of *S. coelicolor afsR* along with its native promoter for insertion into pSS766 with a C-terminal 3x FLAG tag | AAAACGCTCACTGGTACCGCCTCCCC CGCCGCACA |
| SS1912 | Amplification of *S. coelicolor afsR* or afsR$^{GV}$ along with its native promoter for insertion into pSS766 | AGGGCCAGCGCCTCCTCGGG |
| SS1913 | Amplification of *S. coelicolor afsR* or afsR$^{GV}$ along with its native promoter for insertion into pSS766 | GGAGGCGCTGGCCCTCTTCACG |
| SS1914 | Amplification of *S. coelicolor afsR* or *afsR*$^{GV}$ along with its native promoter for insertion into pSS766 | CGTGGTCCTTGTAGTCCTCGAGCCGC GCCACACTGCGAGCCAGCAACGC |
| afsS-BT-FW | Biotinylated forward sequence relating to AfsR binding site in *afsS* promoter for bio-layer interferometry experiments. | GTAGCCGGAGCGTTCAGCGTTCGTTT ATCTCCCCCTGGCACT |
| afsS-Rev | Reverse sequence relating to AfsR binding site in *afsS* promoter for bio-layer interferometry experiments. | AGTGCCAGGGGGAGATAAACGAACGC TGAACGCTCCGGCTAC |
| wblH-BT-Fw | Biotinylated forward sequence relating to AfsR binding site in *wblH* promoter for bio-layer interferometry experiments. | GTTGATACCGCGTTGAGCGAACGTTT TTCGCTTCCCGGCAGG |
| wblH-Rev | Reverse sequence relating to AfsR binding site in *wblH* promoter for bio-layer interferometry experiments. | CCTGCCGGGAAGCGAAAAACGTTCGC TCAACGCGGTATCAAC |
| Scram-BT-Fw | Biotinylated scrambled forward sequence for use as a control in bio-layer interferometry. | TTTCTGGTCGACTGAGCCAAAACTGT GTGTGCTCCCGGTGCG |
| Scram-wblH-Rev | Biotinylated scrambled reverse sequence for use as a control in bio-layer interferometry. | CGCACCGGGAGCACACACAGTTTTGG CTCAGTCGACCAGAAA |

### Datasets

**Supplementary Data 1:** Total number of AfsR, AfsS and WblH orthologs identified by reciprocal BLAST analysis.

**Supplementary Data 2:** RNA-seq data.

**Supplementary Data 3:** Statistically significant ChIP-seq enrichment sites.

**Supplementary Data 4:** Total number of identified AfsR-like proteins.

**Supplementary Data 5:** AfsR-like protein BGC association.

**Supplementary Data 6:** AntiSMASH database BGC annotations of available genomes encoding AfsR-like regulators.

**Supplementary Data 7:** Output of comparative metabolomic analysis of *S. coelicolor.*

**Supplementary Data 8:** Mass list used for untargeted metabolomics analysis of *S. coelicolor* samples.

**Supplementary Data 9:** Output of comparative metabolomic analysis of *S. peucetius*
